# Structures of full-length glycoprotein hormone receptor signaling complexes

**DOI:** 10.1101/2021.08.03.454894

**Authors:** Jia Duan, Peiyu Xu, Xi Cheng, Chunyou Mao, Tristan Croll, Xinheng He, Jingjing Shi, Xiaodong Luan, Wanchao Yin, Erli You, Qiufeng Liu, Shuyang Zhang, Hualiang Jiang, Yan Zhang, Yi Jiang, H. Eric Xu

## Abstract

Luteinizing hormone (LH) and chorionic gonadotropin (CG) are members of the glycoprotein hormone family essential to human reproduction and are important therapeutic drugs. They activate the same G protein-coupled receptor, LHCGR, by binding to the large extracellular domain (ECD). Here we report four cryo-EM structures of LHCGR, two wildtype receptor structures in the inactive and active states, and two constitutively active mutated receptor structures. The active structures are bound to CG and Gs heterotrimer, with one of the structure also containing the allosteric agonist, Org43553. The structures reveal a distinct ‘push and pull” mechanism of receptor activation, in which the ECD is pushed by the bound hormone and pulled by the extended hinge loop next to the transmembrane domain (TMD). A highly conserved 10-residue fragment (P10) from the hinge C-terminal loop at the ECD-TMD interface functions as a tethered agonist to induce conformational changes in TMD and G-protein coupling. Org43553 binds to a TMD pocket and interacts directly with P10 that further stabilizes the receptor in the active conformation. Together, these structures provide a common model for understanding glycoprotein hormone signal transduction and dysfunction, and inspire the search for clinically suitable small molecular compounds to treat endocrine diseases.

---

Luteinizing hormone (LH) and chorionic gonadotropin (CG) are two related hormones that play critical roles in sex development and human reproduction^1, 2^. LH is produced by the pituitary gland and is key to follicle maturation, steroidogenesis, and ovulation. CG is produced by the placenta that is essential to protect the human conceptus during early pregnancy ^1, 3, 4^. Both LH and CG are therapeutic drugs for reproductive and sex development disorders^5, 6^. LH and CG belong to the glycoprotein hormone family, which also includes follicle-stimulating hormone (FSH) and thyroid-stimulating hormone (TSH). These hormones are heterodimers of cysteine-knot proteins, which share a common α-subunit and a unique β-subunit that determines hormone specificity^7–9^. Crystal structures of CG and FSH have been solved, which reveal a similar architecture of an elongated fold for both α- and β-subunits^10, 11^.

The functions of glycoprotein hormones are mediated through a family of closely related G protein-coupled receptors (GPCRs) that belong to class A GPCRs^12^. Members of the glycoprotein hormone family display exquisite selectivity toward their receptors, which contain a large extracellular domain (ECD) of leucine-rich-repeats (LRRs) and a hinge region, followed by a rhodopsin-type seven-transmembrane-helix domain (TMD, Fig.1a). The glycoprotein hormone receptors are also related to leucine-rich-repeat GPCRs (LGRs), which have been proposed as receptors for several cysteine-knot proteins, including R-spondins and norrin^13–16^. These receptors constitute a subfamily branch of class A GPCRs, in which no full-length structure is available. The only available structures are ECD fragments from FSHR, TSHR, LGR4 and LGR5^17–23^. The structures of the FSHR ECD bound to FSH reveal that the hinge region is integrated with the LRRs to form a completed and modular domain of ECD^18^, where FSH is bound to the distal N-terminal regions of ECD with both α- and β-subunits contacting the ECD^17, 18^. The ECD binding by the unique β-subunit provides the basis for the specificity of hormone recognition, in which the binding mode is conserved across glycoprotein hormones^17, 18^.

**Figure 1.**
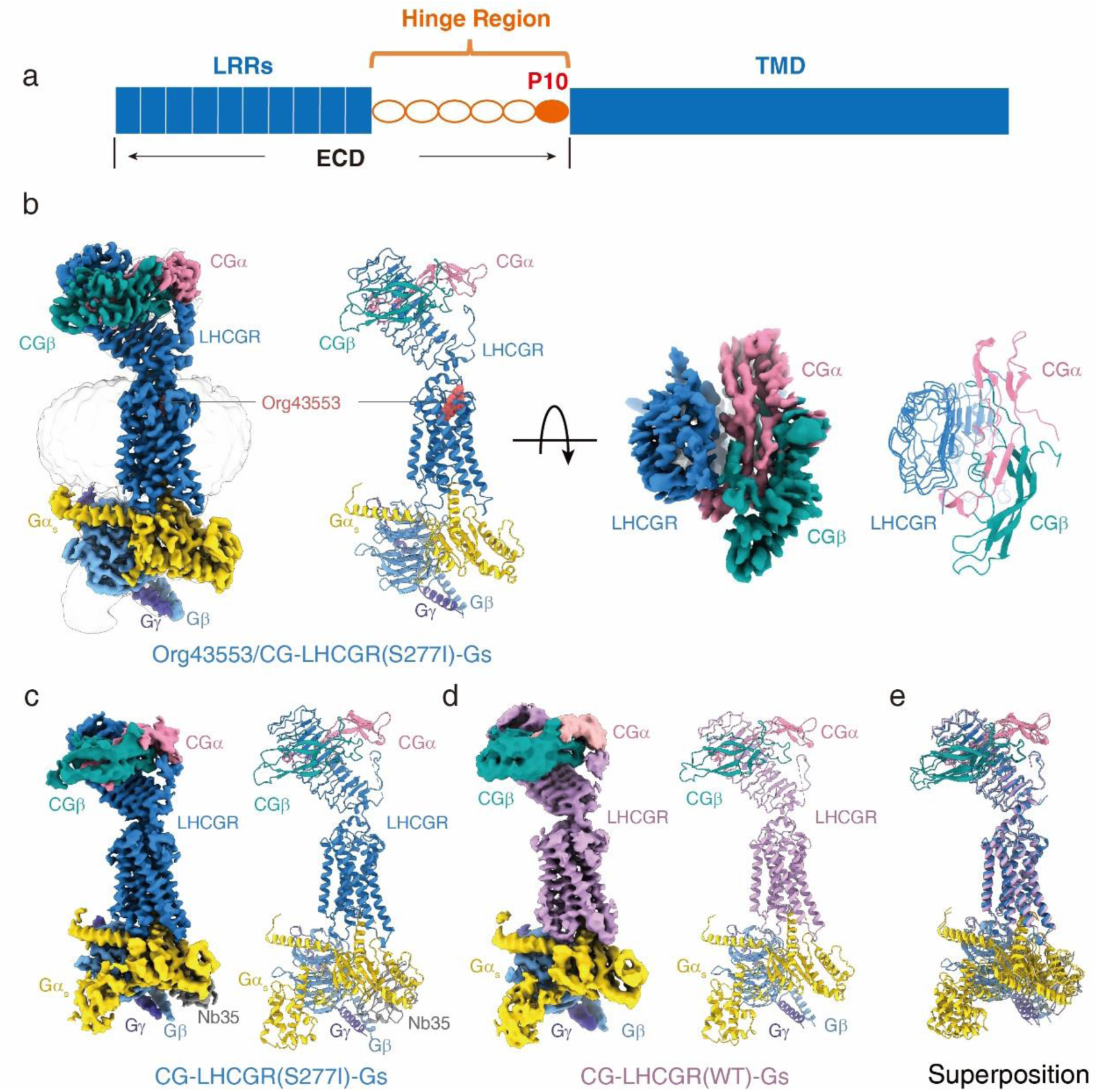
Cryo-EM structure of the CG-LHCGR-Gs complexes. **a**, Schematic diagram of the structure composition of glycoprotein hormone receptors. **b**, Cryo-EM density (left panel) and ribbon presentation (right panel) of the Org43553/CG-LHCGR(S277I)-Gs complex. CGα, pink; CGβ, green; LHCGR, blue; Gα, yellow; Gβ, light blue; Gγ, slate blue; Org43553, red; Nb35, grey. **c**, Cryo-EM density (left panel) and ribbon presentation (right panel) of the CG-LHCGR(S277I)-Gs complex. **d**, Cryo-EM density (left panel) and ribbon presentation (right panel) of the CG-LHCGR(WT)-Gs complex. LHCGR, purple. **e**, Superposition of CG-LHCGR(S277I)-Gs complex with CG-LHCGR(WT)-Gs complex.

However, it remains unclear how the binding of glycoprotein hormones at the distal regions of ECD transmits the binding signal across the receptor TMD to the intracellular G-protein heterotrimer. In addition, there have been great efforts in developing small molecule agonists to replace glycoprotein hormones in clinical applications^24, 25^. Org43553 is a small molecule allosteric agonist of LHCGR in phase 1 clinical trial for replacement of LH and CG, but its mechanism of action remains unknown (https://www.cortellis.com/drug discovery). To address these questions, we determined two structures of wildtype LHCGR in both inactive and active states and two structures of constitutively active mutated receptor with one structure also containing the allosteric agonist, Org43553.

## Assembly and structure determination of the CG-bound LHCGR-Gs complexes

It has been extremely challenging to express the full-length glycoprotein hormone receptors and assemble a fully active hormone bound receptor-Gs complex. We expressed human CG with LHCGR and three subunits of Gs heterotrimer, but failed to assemble the active CG-LHCGR-Gs complex. To overcome these difficulties, we used native human CG from urinary sources. In addition, we introduced a constitutively active mutation, S277I, to enhance the assembly of the CG-LHCGR-Gs complex^26^. Incubation of CG with membranes from cells co-expressing LHCGR and Gs heterotrimer in the presence of Nb35, which stabilizes the receptor-Gs complex^27^, allowed efficient assembly of the CG-LHCGR-Gs complex, which was purified to homogeneity for structural studies (Extended data Fig.1a). Org43553 was added to further stabilize the CG-LHCGR-Gs complex (Extended data Fig.1b).

The structure of the CG-LHCGR-Gs complex alone or bound to Org43553 was determined by single-particle cryo-EM to the resolution of 3.9 Å and 3.18 Å, respectively (Fig.1b-c, Extended data Fig.1c-f). For the Org43553/CG-LHCGR-Gs complex, the EM map was clear to position all components of the complex, including two subunits of CG, LHCGR, the three Gs subunits, and Nb35 (Fig.1b, Extended data Fig.1d, f-g). Residues 52-644 of LHCGR, 5-92 of CG α-subunit, and 2-111 of CG β-subunit were included in the final model. We also determined a structure of wild type LHCGR in complex with CG and Gs protein at 4.3 Å resolution (Fig.1d, Extended data Fig.2a, c, e), and the overall structure of this complex is largely overlapped with the S277I mutated receptor complex (Fig.1e). The data and structure statistics are summarized in extended data Table 1. Because the structure of the S277I mutated LHCGR in complex with CG, Gs protein and Org43553 has the best resolution, the structural analysis will be based on this structure unless indicated otherwise.

## Basis for hormone recognition of CG by LHCGR

In the CG-LHCGR complexes, both CG subunits adopt a similar elongated fold of four β-strands that are stabilized by the core cysteine-knot scaffold (Extended data Fig.3a). The C-terminal segment of the β-subunit serves as a “seat belt” that encircles the α-subunit (Extended data Fig.3b). In the active structures, CG binds to the top end of the ECD, far from the TMD (Fig.1b-d). The LHCGR ECD contains 11 irregular LRRs that form a slightly curved tube, where CG binds to the concave inner surface of the LRRs in a hand-clasp fashion (Fig.2a-b, Extended data Fig.3c). The hinge region contains LRR10, an α-helix (termed hinge helix), and LRR11, which together with LRR1-9 form the complete ECD, resembling the complete ECD structure of FSHR^18^. Between the hinge helix and LRR11 is an extended hinge loop (residues 284-340), of which only the C-terminal portion is visible in the structure; part of this region appears to interact with the interface formed by CG α- and β-subunits (Fig.2a-b). The surface of CG contains several discrete positively charged patches, forming complementary electrostatic interactions with negatively charged patches on the surface of the LHCGR ECD (Fig.2c).

**Figure 2.**
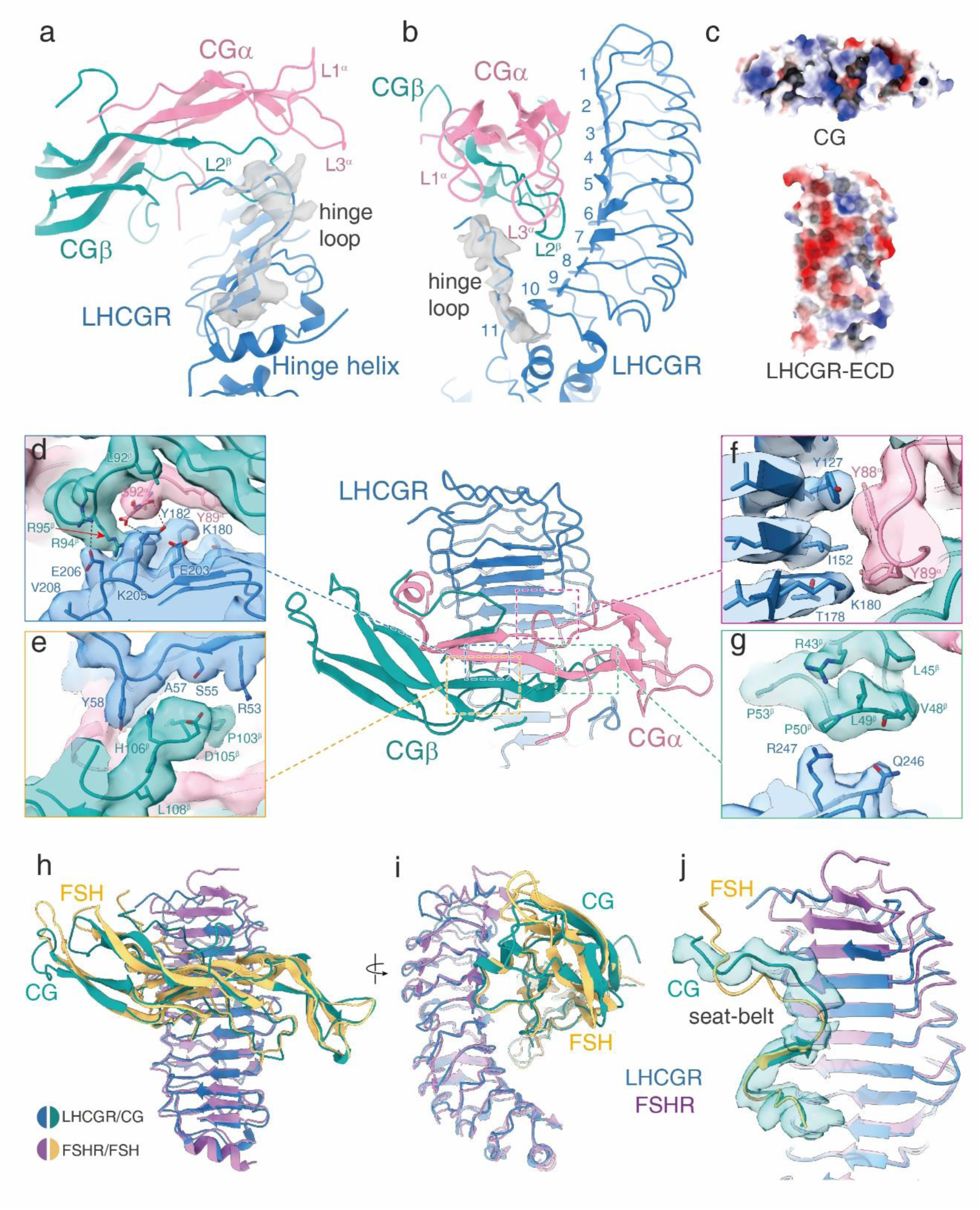
Structural basis of CG/LHCGR ECD interaction. **a,b** The interaction interface of the LHCGR hinge loop with CGα and CGβ subunits; The density map of the extended C-terminus of hinge loop is shown in grey. **c**, Surface charge distribution of CG and LHCGR-ECD. **d-g**, The interaction interface of LHCGR-ECD with CGα (pink) and CGβ (green). **h-j,** Structural comparison of the CG-LHCGR complex with the FSH-FSHR complex (PDB code: 1XWD). The overall binding mode of the two hormones to the receptor ECDs (**h, i**); The “seat-belt” fragment in the C-terminus of CG β-subunit is indicated; Structural comparison of the interface formed by “seat-belt” fragments and the receptor ECDs (**j**). Residues’ side chains are displayed as sticks. Polar interactions are shown in black dashed lines.

The interactions of CG with LHCGR are mediated by both CG subunits (Fig.2d-g), and are summarized in extended data Figure 4 and Table 2. Binding of the CG α-subunit is primarily mediated by Y88^α^, Y89^α^, and S92^α^, which form packing interactions with Y127, I152, K180 and Y182 from LRR4-6 of LHCGR (Fig. 2d, f, Extended data Fig.4a, c). Binding of the CG β-subunit is primarily mediated by its C-terminal residues (92 to 106) (Fig. 2d-e, Extended data Fig. 4a-b), which form close interactions with R53, S55, A57 and Y58 from LRR1 (Fig.2e), and with E206 from LRR7(Fig. 2d). Additional interactions by the CG β-subunit are seen in residues V46^β^ and Q48^β^, which form interactions with Q246 and R247 from LRR10 (Fig.2g). Compared to free CG structures, the receptor-bound CG displays conformational changes in four segments (β-subunit “seat-belt”, βL2, βL3, and αL3), three of which are near the interface with the receptor (Extended data Fig.3d).

Structural comparison between the CG-LHCGR and the FSH-FSHR complexes reveals the basis for the specificity of hormone recognition (Fig.2h-j, Extended data Fig.3e). Although the overall binding modes of both hormones to the receptors are very similar^17^, particularly by the common α-subunits, the detailed interactions by β-subunits are actually quite different. Specifically, the C-terminal “seat-belt” segments of β-subunits adopt very different conformations, in which the Cα atom of P107^β^ in CG is 6.4 Å apart from its counterpart in P101^β^ of FSH (Fig.2j, Extended data Fig.3e). Neighboring P101^β^ in FSH are L99^β^ and Y103^β^, which are known to form a hydrophobic pocket to specifically accommodate the side chain of L55 of FSHR^17^. The corresponding residue in LHCGR is the much bulkier Y58 (Fig. 2e), which would have a severe clash with L99^β^ of FSH, suggesting that this is a key site of structural discrimination in determining specificity. The corresponding C-terminal “seat-belt” in the LH β-subunit is conserved in CG, thus allowing LHCGR to adopt LH binding (Extended data Fig.4b). Additional specificity lies in R94^β^ in the CG β-subunit, which forms a salt bridge with E206 of LHCGR(Fig.2h). The corresponding residues are S89^β^ in FSH and D202 of FSHR. On the other hand, FSH has a specific D90^β^, which forms a salt bridge with K179 of FSHR. These two corresponding residues are not present in CG and LHCGR, further illustrating the basis of hormone recognition specificity between the two signaling systems of CG-LHCGR and FSH-FSHR.

## Basis for hormone-induced receptor activation

LHCGR has a low level of basal activation in the absence of CG^28^. In order to reveal the mechanism of hormone-induced receptor activation, we attempted to determine both the active and inactive structures of LHCGR and compare these structures to reveal the hormone-induced conformational changes. After exhaustive efforts, we were able to solve the inactive wildtype LHCGR structure to 3.8 Å, which was sufficient to place the ECD and the TMD (Fig.3a-b, Extended data Fig.2b, d, f, g). The ECD in the inactive structure is close titled toward the membrane layer, which is in contrast to the active LHCGR complex, where the ECD is nearly perpendicular to the membrane layer (Figure 3b-c). It appears that the binding of CG induces an upward rotation of this domain by about 45 degrees (Fig. 3c-d). The inactive LHCGR TMD is very much similar to the inactive conformation of class A GPCRs, including rhodopsin^29^ and β_2_ adrenergic receptor (β_2_AR)^30^ (Extended data Fig.3f), particularly at the C-terminal end of TM6, supporting the contention that the inactive LHCGR is indeed in the inactive state. Comparison with the CG-bound LHCGR-Gs complex reveals that the C-terminal end of TM6 undergoes an outward movement upon hormone binding by as much as 12.3 Å as measured at the Cα atom of residue D564^6.30^(superscripts refer to Ballesteros-Weinstein numbering^31^)(Fig.3e). Accompanying TM6 outward movement are 2 Å outward movement of TM5 and 3.6 Å inward movement of TM7 (Fig. 3e), in agreement with the general paradigm of class A GPCR activation.

**Figure 3.**
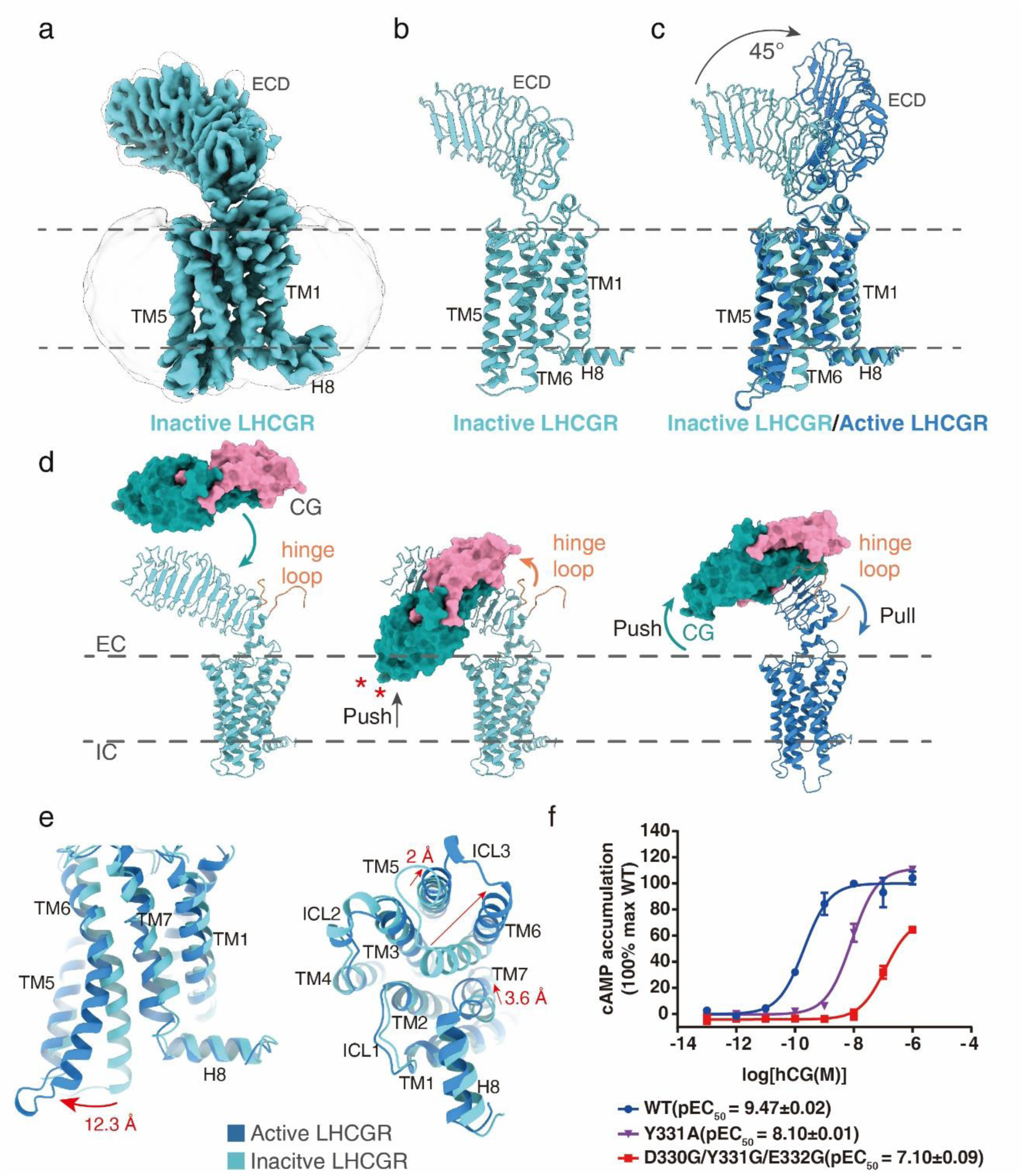
Basis for hormone-induced receptor activation. **a,b** EM density and ribbon presentation of inactive LHCGR. **c**, Comparison of inactive and active LHCGRs. The ECD in active LHCGR undergoes a 45°upward movement relative to inactive LHCGR. **d**, The putative activation model of LHCGR. The binding of CG to the inactive LHCGR would cause a clash of the distal region of CG with the membrane layer, pushing the rotation of the ECD-CG complex into the upward-pointing configuration. The C-terminus of the extended hinge loop may interact with CG, further pulling the CG-ECD complex in the upward position. **e**, Conformational comparison of the TMD cytoplasmic part of inactive and active LHCGR. **f**, Concentration-response curves for point mutants in hinge loop. Experiments were performed in triplicate, and the representative concentration-response curves were presented.

The key question is how the binding of CG at the distal ECD induces the activation of the LHCGR TMD. As illustrated above (Fig.3d), the binding of CG to the inactive LHCGR would cause a clash of the distal region of CG with the membrane layer, which would push the rotation of the CG-ECD complex into the upward-pointing configuration. On the other hand, we observed a trace of poorly-resolved EM density corresponding to the C-terminal portion of the extended hinge loop that interacts with the far right tip of CG (Fig.2a-b), in a manner similar to the FSH-FSHR ECD complex^18^. We propose that this CG interaction by the hinge loop serves to pull the CG-ECD complex into the upward position. Within the middle of the hinge loop is a sulfonated tyrosine (Y331) surrounded by negatively charged residues^32^, which have been shown to interact with a positively charged patch at the CG surface (Extended data Fig.3g). Mutations of these residues to uncharged residues (Y331A and D330G/Y331G/E332G mutants) result in reduced activation of the receptor (Fig.3f, Extended data Table 2). Together, these results illustrate a push and pull model of LHCGR activation by CG binding.

In the CG-LHCGR-Gs complex, there are extensive ECD-TMD interactions stabilized by the upward movement of ECD induced by CG binding (Fig.4a-c, Extended data Table 2). Two major ECD-TMD interfaces are observed in the structure. The first interface is mediated by the ECD hinge helix, which is packed closely against the ECL1, which also adopts a two and half-turn helix (Fig.4a). The constitutively active S277I mutation is located at the N-terminus of the hinge helix and this mutation increases the ECD-TMD binding interface. Specifically, S277I forms additional hydrophobic interactions with A430 and I431 from the ECL1 helix, thus stabilizing the ECD-TMD interactions. Consistently, A430D, S277K, and S277R mutations, which are likely to affect the ECD-TMD interface and decreased activities of the receptor (Extended data Fig.3h and Table 2). In the wildtype receptor structure, the ECD hinge helix adopts a similar conformation with S277 at the same position(Fig.1e). However, S277 would not be able to form the same hydrophobic interactions as S277I, instead S277 is in a position to form a possible “hydrogen bond” with N351 from the conserved P10 region (residue 350-359) (Extended data Fig.3i), which has been posposed as the tethered agonist for LHCGR and TSHR^33^.

**Figure 4.**
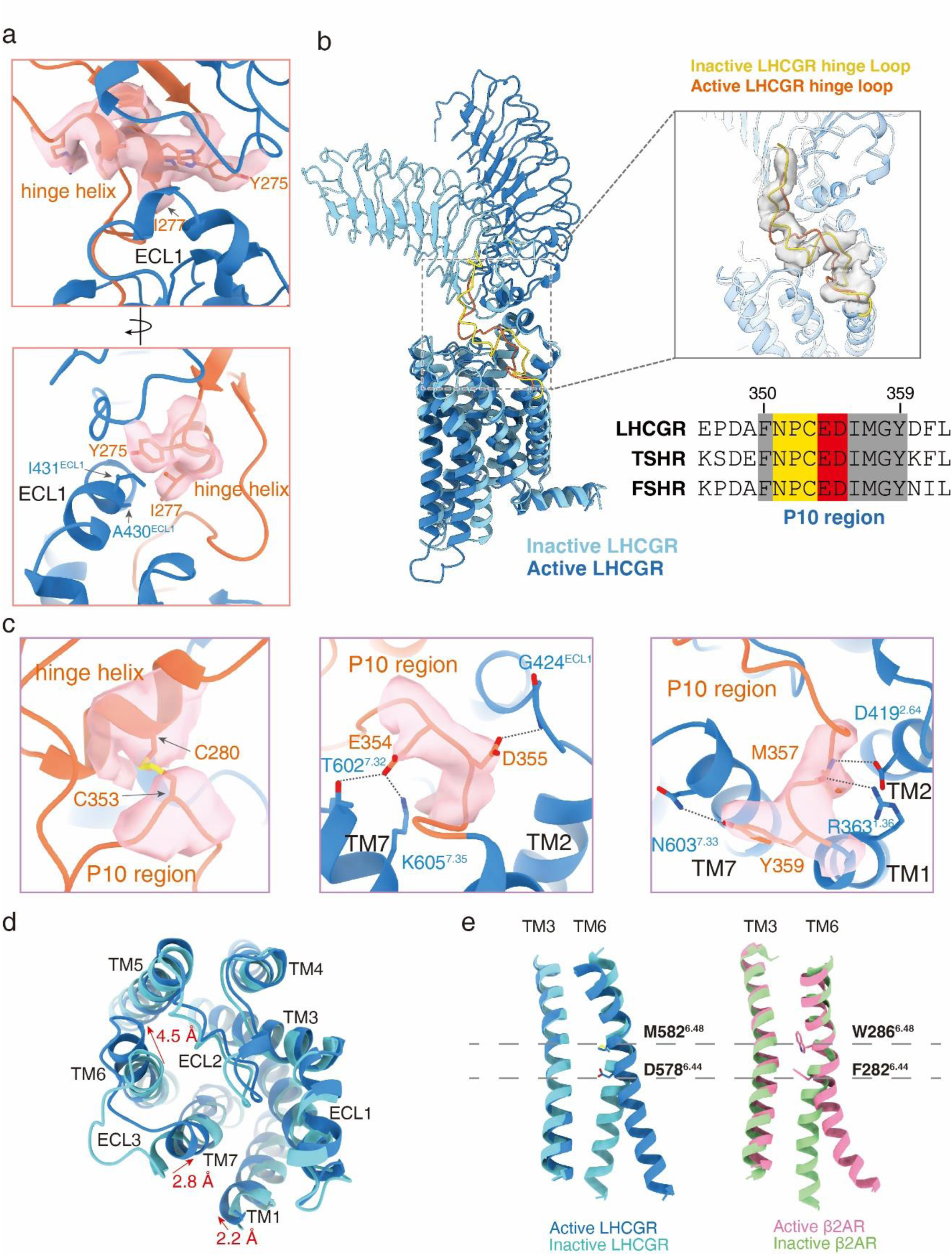
Interactions between LHCGR ECD and TMD. **a**, The interface of ECL1 and hinge helix. **b**, Structure comparison of P10 region in the active and inactive LHCGR. And sequence alignment of the hinge C-terminus fragment among LHCGR, TSHR and FSHR. The P10 region was conserved among three receptors. **c**, Detail interactions between TMD and the P10 region. The EM densities of side chains are indicated in light pink. **d**, Conformational comparison of the TMD extraplasmic part of inactive and active LHCGR. **e**, Structural representation of the TM3 and TM6 in LHCGR and β_2_AR. The residues M582^6.48^ and D578^6.44^ at the TM6 helical kink in LHCGR were shown as sticks, homologous residues to the toggle switch of W286^6.48^ and the PIF motif residue F282^6.44^ in β_2_AR were also as sticks.

The second ECD-TMD interface is formed by the hinge C-terminal P10 regrion, which is packed at the center-top of the TMD bundle and forms extensive interactions with TM1, TM2, and TM7 as well as all three ECLs (Fig.4b-c). In the inactive structure, this region adopts a different conformation from the active structure (Fig. 4b), suggesting that receptor activation involved conformational rearrangement of the P10 region. The interactions of P10 region with the TMD observed in our structures are well correlated with previous structure prediction and mutational studies^33, 34^. This fragment is one of the most conserved regions in the glycoprotein hormone receptors(Fig.4b), suggesting a highly conserved activation mechanism across glycoprotein hormone receptors. Additional differences in the active structure are seen in the extracellular ends of TM6 and TM7, which move by 3-5 Å from the inactive structure (Fig.4d). These movements lead to a helical kink at M582^6.48^ and D578^6.44^, analgous residue to the toggle switch of W286^6.48^ and the PIF motif residue F282^6.44^ in β_2_AR, which are two key components in the activation of class A GPCRs. The residue types of M582^6.48^ and D578^6.44^ are conserved in glycoprotein hormone receptors but are different from most class A GPCRs (Extended data Table 3), suggesting that glycoprotein hormone receptors have evolved a unique activation mechanism from other class A GPCRs.

The overall structure of the CG-LHCGR-Gs complex is organized into an integrated and interconnected complex rather than disconnected modular domain structures between ECD and TMD. In molecular dynamic simulations, the distances from ECD to membrane and the tilting angles of ECD were maintained throughout the course of simulation in the presence of CG but not in the absence of CG, further reinforcing the notion that the binding of CG to ECD serves to stabilize and rigidify the ECD-TMD interface (Extended data Fig.5a-b). In addition, the CG-LHCGR-Gs complex structure is in a monomeric state, which is consistent with all class A GPCR-Gs complex structures solved to date. Although the crystal structure of the FSH-FSHR-ECD complex has been determined in a dimeric or trimeric state^17, 18^, the rigid CG-LHCGR-Gs complex would prevent receptor dimerization or trimerization in the presence of glycoprotein hormones and G proteins in the same membrane layer from the same cell (Extended Fig.5c-d).

## Basis for LHCGR regulation by allosteric agonist Org43553

Org43553 is a drug candidate that has been reported as an allosteric agonist of LHCGR^35^. The EM map reveals that Org43553 inserts into a deep pocket formed by the top half of the TMD (Fig.5a-b). Org43553 occupies the top portion of the pocket with morpholine ring exposed to solvent and the ECD-TMD interface. Molecular dynamic simulations indicate that the binding mode of Org43553 is relatively stable within the binding pocket during the course of simulations (Extended Fig.6a-d). The binding pocket is made up by residues from TM3, TM5, TM6, and TM7, as well as residues from ECL2, ECL3, and the hinge C-terminus (Fig.5d). Org43553 engages LHCGR predominantly by hydrophobic interactions (Fig.5d-e). Mutations of A589W and I585F, which protrude into the Org43553-binding pocket, reduced the ability of Org43553 to activate LHCGR (Fig.5c), supporting the binding mode of Org43553 observed in the structure.

**Figure 5.**
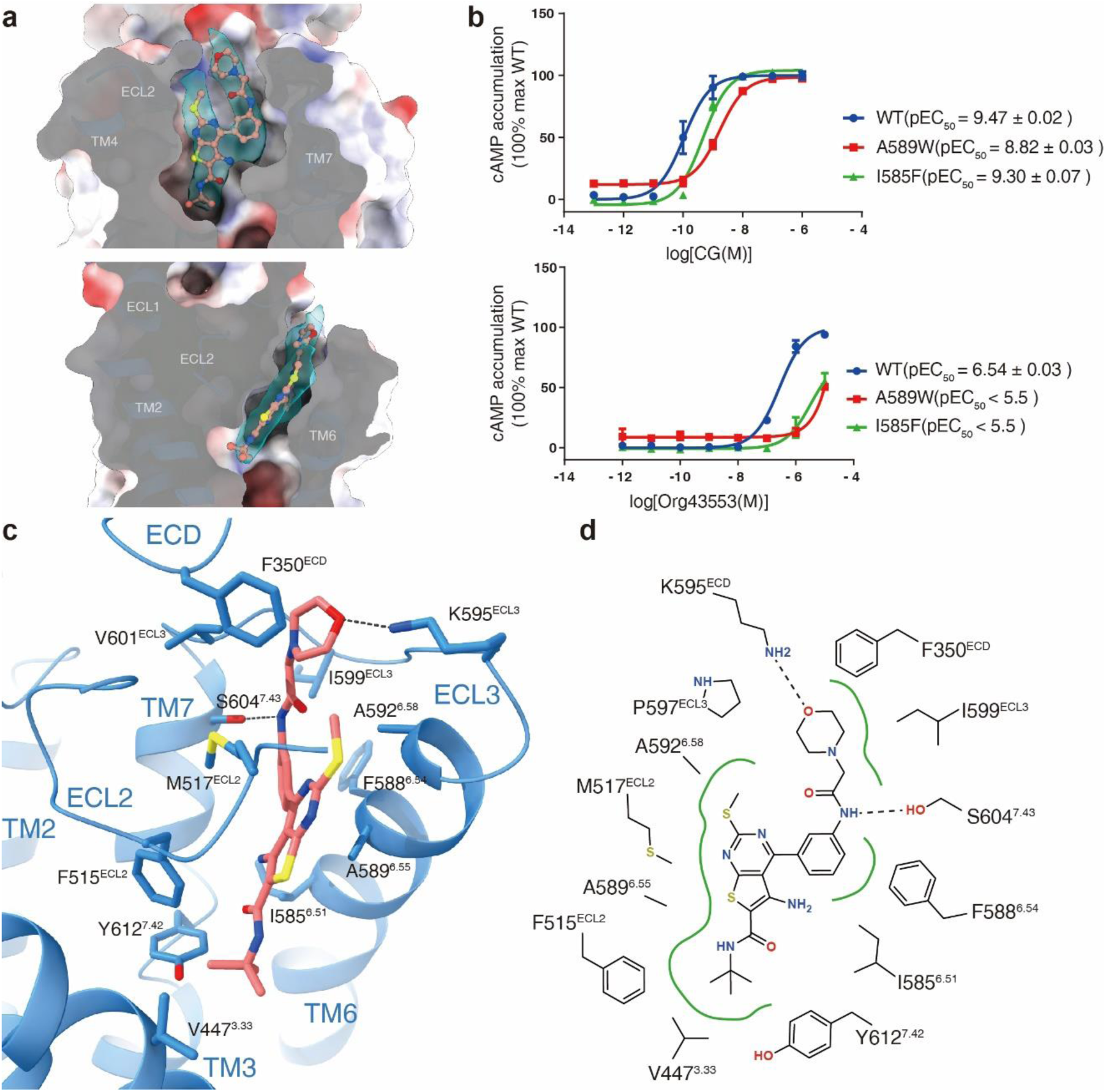
Basis for LHCGR regulation by Org43553. **a,** The EM density presentation of Org43553 in full-length LHCGR. **b,** Concentration-response curves to demonstrate the effects of A589W and I585F mutations on CG and Org43553-induced LHCGR activation. Experiments were performed in triplicate, and the representative concentration-response curves were presented. **c**, Detail interactions between LHCGR and Org43553. Side chains of residues in the Org43553-binding pocket are shown as sticks. **d**, Schematic representation of Org43553-binding interactions. Hydrogen bonds are shown as black dashed lines. Hydrophobic interactions and amino acids are shown in green.

The position of Org43553 is far away from the binding site of CG at the distal end of ECD. Compared to other class A GPCRs, the location of Org43553 is similar to other orthosteric agonists; in particular, its *tert*-butylamine group overlaps with the space occupied by the bottom groups of other class A GPCR agonists (Extended data Fig.6d). These structural observations suggest that Org43553 is a direct TMD activator of LHCGR in a manner similar to orthosteric agonists of class A GPCRs. Consistently, Org43553 can activate LHCGR in the absence of CG (Fig. 5c)^35^.

Structural comparision of the CG-LHCGR complexes with and without Org43553 reveals conformational changes in LHCGR upon the binding of Org43553 (Extended data Fig.7). The morpholine ring of Org43553 interacts with the F350 of the receptor, pushing the hinge C-terminus to produce a 1 Å shift, which further lead to a movement of about 2 Å in the hinge helix (Extended data Fig.7a). Additional changes are small 1-2 Å movements in the extracellular ends of TM helices (Extended data Fig.7b).

Correspnding, we also observed small changes in the cytoplamic ends of TM helices but no obvious changes in the CG/ECD interface between the two structures(Extended data Fig.7a,c).

## Basis for LHCGR coupling with Gs protein

The overall structure of the LHCGR-Gs complex exhibits the common mode of GPCR-Gs coupling, including the insertion of the C-terminal α5 helix of Gαs into the cytoplasmic cavity of the transmembrane helix bundle. Y391 near the C-terminal end of α5 helix forms a stacking interaction with R464^3.50^, which is from the highly conserved DRY motif of class A GPCRs^27, 36–38^ (Extended data Fig.8d). Asides from these common features, LHCGR displays several distinct features in its interface with the Gs heterotrimer. When compared to the structure of the β_2_AR-Gs complex, the overall Gs complex is rotated by 10-15 degrees counter-clockwise, as observed for the αN helix and the C-terminal α5 helix of Gαs (Extended data Fig.8a-c). The C-terminal α5 helix of Gαs inserts 3 Å (measured at the Cα atom of residue Y391) deeper into the cytoplasmic TMD pocket, which allows the direct interaction of the α5 helix C-terminus with the N-terminus of helix 8 and the turn of TM7-helix 8 (Extended data Fig.8c-d). In addition, ICL2 did not adopt a helical conformation like ICL2 in the β_2_AR-Gs complex (Extended data Fig.8e).

Together, these structural comparisons reveal the common and unique mechanisms of Gs coupling by LHCGR and other GPCRs.

## Disease-associated mutations in LHCGR

Missense mutations in LHCGR have been associated with a number of diseases such as infertility, and male-limited precocious puberty^2,26, 39, 40^. The impact of most mutations can be rationalized based on our structure. Based on the location in the LHCGR structure, these mutations can be classified into inactive and constitutively active mutations (Extended data Fig.9 and Fig.10). The inactive mutations in the ECD are either located in the CG-LHCGR interface or in the LRR hydrophobic core. Other inactive mutations are spread accorss the TMD with many mutations forming the TMD core. Most constitutively active mutations are located within TMD, with seven mutations in the cytoplasmic half of TM6, including mutations at D578^6.44^, which located at the kink of TM6. Only one constitutively active mutation was found respectively from TM3, TM5, and TM7, and all these three mutations are in close proximity to D578^6.44^. Together, these mutations are likely destabilize TM6 from it inactive conformation. S277N is the only constitutively active mutation naturally occurring in a patient^40^, mutations at S277 is the only single constitutively active mutations outside of TMD, which located at the ECD-TMD interface that stabilize ECD-TMD interaction as discussed earlier. Because of the conservation in the glycoprotein hormone receptors, the LHCGR structures are likely to serve as models for understanding endocrine disease associated mutations occurring in this family of receptors.

In summary, we have overcome technical difficulties to determine multiple structures of the full-length LHCGR, in the inactive and hormone-induced active states. Glycoprotein hormones are well known for their important physiological functions and for their specificity in the activation of their receptors. Our structures have addressed long-standing questions regarding the structural architecture of the glycoprotein hormone receptor signaling complex and the mechanism of hormone-induced receptor activation. Specifically, our structures reveal a push and pull model of CG binding to the LHCGR ECD that stabilizes the ECD-TMD interactions, which ultimately leads to the receptor activation. Our structures also reveal the binding site and the docking mode of Org43553 within the TMD. Importantly, our active CG-LHCGR-Gs complex is in a monomeric state like most class A GPCR-G protein complexes, rather than a dimeric or trimeric state because the rigid ECD-TMD interface observed in our structure would prevent the oligomerization previously proposed for this complex^17, 18, 41^. Given the highly conserved sequences in glycoprotein hormones and their receptors, the structural mechanisms of hormone-induced activation are likely conserved across this family of hormone receptors. In addition, the structure of the Org43553-bound complex provides crucial information for the drug design of small molecule agonists with aims to treat diverse disorders related to glycoprotein hormones.

## Method

### Constructs

Human LHCGR (residues 27-699) was cloned with an N-terminal FLAG and C-terminal His8 tags using homologous recombination (CloneExpress One Step Cloning Kit, Vazyme). Additional mutation S277I was designed to form CG-LHCGR(S277I)-Gs complexes. The native signal peptide was replaced with the haemagglutinin (HA) to increase protein expression. A dominant-negative bovine Gαs construct was generated based on mini-Gs^36^. The N-terminal 1-18 amino acids and α-helical domain of Gαs were replaced by human Gαi1, thus providing binding sites for ScFV16 and Fab-G50, respectively^42, 43^. Additionally, three mutations (G226A, A366S and L272D) were also incorporated by site-directed mutagenesis to decrease the affinity of nucleotide-binding and increase the stability of Gαβγ complex^44^. Rat Gβ1 was cloned with an N-terminal His6 tag. All the three G-protein components, including bovine Gγ2, were cloned into a pFastBac vector, respectively.

### Expression and purification of Nb35

Nanobody-35 (Nb35) with a C-terminal His6 tag, was expressed and purified as previously described^27^. Nb35 was purified by nickel affinity chromatography (Ni Smart Beads 6FF, SMART Lifesciences), followed by size-exclusion chromatography using a HiLoad 16/600 Superdex 75 column and finally spin concentrated to 5 mg/ml.

### Complexes expression and purification

LHCGR, Gαs, Gβ1 and Gγ2 were co-expressed in *Sf9* insect cells (Invitrogen) using the Bac-to-Bac baculovirus expression system (ThermoFisher). Cell pellets were thawed and lysed in 20 mM HEPES (Sangon), pH 7.4, 100 mM NaCl, 10% glycerol, 5 mM MgCl_2_ and 5 mM CaCl_2_ supplemented with Protease Inhibitor Cocktail, EDTA-Free (Roche). The CG-LHCGR-Gs complex was formed in membranes by the addition of 1 μM CG (Yuke chemical), 10 μg/mL Nb35 and 25 mU/mL apyrase. The suspension was incubated for 1.5 h at room temperature. The membrane was then solubilized using 0.5% (w/v) n-dodecyl β-D-maltoside (DDM, Anatrace), 0.1% (w/v) cholesterol hemisuccinate (CHS, Anatrace) and 0.1%(w/v) sodium cholate for 2 h at 4℃. The supernatant was collected by centrifugation at 80,000 × g for 40 min and then incubated with M1 anti-Flag affinity resin for 2 h at 4℃. After batch binding, the resin was loaded into a plastic gravity flow column and washed with 10 column volumes of 20 mM HEPES, pH 7.4, 100 mM NaCl, 10% glycerol, 2 mM MgCl_2_, 2 mM CaCl_2_, 0.01% (w/v) DDM, 0.002%(w/v) CHS, and 0.002%(w/v) sodium cholate, 0.1 μM CG and further washed with 10 column volumes of same buffer plus 0.1%(w/v) digitonin, and finally eluted using 0.2 mg/mL Flag peptide. The complex was then concentrated using an Amicon Ultra Centrifugal Filter (MWCO 100 kDa) and injected onto a Superdex200 10/300 GL column (GE Healthcare) equilibrated in the buffer containing 20 mM HEPES, pH 7.4, 100 mM NaCl, 2 mM MgCl_2_, 2 mM CaCl_2_, 0.05 (w/v) digitonin, 0.0005% (w/v) sodium cholate, 0.05 μM CG. For Org43553 bound complex, an additional 10 μM Org43553 was added for complex formation and 1 μM Org43553 for the following procedures. The complex fractions were collected and concentrated for electron microscopy experiments, respectively.

### Inactive LHCGR expression and purification

Sf9 insect cells were infected with LHCGR baculovirus and cultured for 48 h at 27 ℃ before collection. Cell pellets were thawed and lysed in 20 mM HEPES, pH 7.4, 100 mM NaCl, 10% glycerol, supplemented with Protease Inhibitor Cocktail, EDTA-Free (Roche). The purification procedures were similar to the CG-LHCGR-Gs complex, except that there was a small molecular antagonist (compound 26)^45^ instead of CG ligand in the buffer.

### cAMP response assay

The full-length LHCGR (27–699) and LHCGR mutants were cloned into pcDNA6.0 vector (Invitrogen) with a FLAG tag at its N-terminus. CHO-K1 cells (ATCC, #CCL-61) were cultured in Ham’s F-12 Nutrient Mix (Gibco) supplemented with 10% (w/v) fetal bovine serum. Cells were maintained at 37 °C in a 5% CO_2_ incubator with 150,000 cells per well in a 12-well plate. Cells were grown overnight and then transfected with 1 μg LHCGR constructs by FuGENE® HD transfection reagent in each well for 24 h. The transfected cells were seeded onto 384-well plates. cAMP accumulation was measured using the LANCE cAMP kit (PerkinElmer) according to the manufacturer’s instructions. Fluorescence signals were then measured at 620 and 665 nm by an Envision multilabel plate reader (PerkinElmer). Data were presented as means ± SEM from at least three independent experiments.

### Detection of surface expression of LHCGR mutants

The cell seeding and transfection follow the same method as cAMP response assay. After 24 h of transfection, cells were washed once with PBS and then detached with 0.2% (w/v) EDTA in PBS. Cells were blocked with PBS containing 5% (w/v) BSA for 15 min at room temperature (RT) before incubating with primary anti-Flag antibody (diluted with PBS containing 5% BSA at a ratio of 1:300, Sigma) for 1 h at RT. Cells were then washed three times with PBS containing 1% (w/v) BSA and then incubated with anti-mouse Alexa-488-conjugated secondary antibody (diluted at a ratio of 1:1000, Invitrogen) at 4 °C in the dark for 1 h. After another three times of wash, cells were harvested, and fluorescence intensity was quantified in a BD Accuri C6 flow cytometer system (BD Biosciences) at excitation 488 nm and emission 519 nm. Approximately 10,000 cellular events per sample were collected and data were normalized to WT.

### Cryo-EM grid preparation and data collection

For the preparation of cryo-EM grids, 3 μL of the purified protein at 12 mg/mL for the CG-LHCGR-Gs complex, 20 mg/mL for the Org43553/CG-LHCGR-Gs complex, 16 mg/mL for the CG-LHCGR(WT)-Gs complex and 8 mg/mL for the inactive LHCGR, were applied onto a glow-discharged holey carbon grid (Quantifoil R1.2/1.3). Grids were plunge-frozen in liquid ethane using Vitrobot Mark IV (Thermo Fischer Scientific). Frozen grids were transferred to liquid nitrogen and stored for data acquisition.

Cryo-EM imaging of the CG-LHCGR-Gs complex, CG-LHCGR(WT)-Gs complex and inactive LHCGR were performed on a Titan Krios at 300 kV in Cryo-Electron Microscopy Research Center, Shanghai Institute of Materia Medica, Chinese Academy of Sciences (Shanghai China), and cryo-EM imaging of the Org43553/CG-LHCGR-Gs complex was performed on a Titan Krios at 300 kV in the Shuimu BioSciences Ltd (Beijing China). A total of 5,393 movies for the CG-LHCGR-Gs complex, 5,327 movies for the CG-LHCGR(WT)-Gs complex and 13,475 movies for the inactive LHCGR protein, were collected with a Gatan K3 Summit direct electron detector with a Gatan energy filter (operated with a slit width of 20 eV) (GIF) at a pixel size of 1.045 Å using the SerialEM software^46^. The micrographs were recorded in counting mode at a dose rate of about 22 e/Å^2^/s with a defocus ranging from -1.2 to -2.2 μm. The total exposure time was 3 s and intermediate frames were recorded in 0.083 s intervals, resulting in a total of 36 frames per micrograph. For the Org43553/CG-LHCGR-Gs complex, a total of 3,410 movies were collected by a Gatan K3 Summit direct electron detector with a Gatan energy filter (operated with a slit width of 20 eV) (GIF) at a pixel size of 1.08 Å using the EPU software. The micrographs were recorded in counting mode at a dose rate of about 18.0 e/Å^2^/s with a defocus ranging from -1.2 to -2.2 μm. The total exposure time was 3.33 s and intermediate frames were recorded in 0.104 s intervals, resulting in a total of 32 frames per micrograph.

### Image processing and map construction

Dose-fractionated image stacks were aligned using MotionCor2.1^47^. Contrast transfer function (CTF) parameters for each micrograph were estimated by Gctf^48^. For the CG-LHCGR-Gs complex, particle selections for 2D and 3D classifications were performed using RELION-3.0^49^. Automated particle picking yielded 1,976,506 particles that were subjected to reference-free 2D classification to discard poorly defined particles, producing 746,223 particles. The initial reference was produced by ab-initio reconstruction for 3D classification, resulting in one well-defined subset with 482,806 particles. The selected subset was performed 3D classification with mask on the LHCGR and CG, resulting two well-defined subsets, which were subsequently subjected to 3D refinement and Bayesian polishing. The final refinement generated a map with an indicated global resolution of 3.9 Å with 355,345 particle projections at a Fourier shell correlation of 0.143. Local resolution was determined using the Resmap^50^ package with half maps as input maps. For the Org43553/CG-LHCGR-Gs complex, automated particle picking using Relion yielded 2,870,315 particles. The particles were imported to CryoSPARC^51^ for 3 rounds of heterogeneous refinement. Two good subsets with 1,028,017 particles were further re-extracted in Relion. The particles were subjected to 2 rounds of 3D classification with a mask on the TMD-Gs region and the complex, respectively, producing one high-quality subset accounting for 413,532 particles. The particles were subsequently subjected to 3D refinement, CTF refinement and Bayesian polishing. The global refinement map shows an indicated resolution of 3.2 Å at a Fourier shell correlation of 0.143. To further improve map quality, the focused 3D refinement on the hCG/ECD and TMD/Gs regions were performed. The focused refinement maps for the TMD/Gs and hCG/ECD regions show the global resolutions of 3.1 Å and 3.3 Å, which were combined using “vop maximum” command in UCSF Chimera^52^. This composite map of the complex was used for subsequent model building.

For the CG-LHCGR(WT)-Gs complex, automated particle picking yielded 6,544,295 particles that were subjected to reference-free 2D classification to discard poorly defined particles. The CG-LHCGR-Gs complex map low-pass filtered to 60 Å was used as the initial reference for 3D classification, resulting in one well-defined subset with 2,914,026 particles. The selected subset was performed additional 3D classification, resulting a well-defined subset, which were subsequently subjected to 3D refinement and Bayesian polishing. The final refinement generated a map with an indicated global resolution of 4.3 Å with 620,536 particle projections at a Fourier shell correlation of 0.143. Local resolution was determined using the Resmap^50^ package with half maps as input maps.

For the inactive LHCGR, image stacks were subjected to beam-induced motion correction using MotionCor2^47^. CTF parameters for non-dose weighted micrographs were determined by Gctf^48^. Cryo-EM data processing was performed using Relion-3.1^49^,CryoSPARC-v2.14^51^, and CisTEM-1.0.0^53^. Automated particle selection using Gaussian blob detection yielded 11,702,165 particles using Relion. The particles were subjected to two rounds of 3D classification to discard poorly defined particles. The good subset with 3,077,024 particles was further imported to CryoSPARC for 2 rounds of heterogeneous refinement. The well-defined subset accounting for 1,067,644 particles was re-extracted in Relion and was subjected to 2 rounds of 3D classification as well, producing one good subset accounting for 311,538 particles. Finally, the particles were imported to CisTEM for a final 3D refinement, which generates a map with an indicated global resolution of 3.8 Å at a Fourier shell correlation of 0.143. Global and local resolution was determined using the Resmap^50^ package with half maps as input maps.

### Model building and refinement

The crystal structure of human CG (PDB code: 1HRP), the structure of rhodopsin (PDB code: 4ZWJ) and the Gs protein model (PDB code: 3SN6) were used as the start for model rebuilding and refinement against the electron microscopy map. The model was docked into the electron microscopy density map using Chimera^52^, followed by iterative manual adjustment and rebuilding in COOT^54^ and ISOLDE^55^. Real space and reciprocal space refinements were performed using Phenix programs^56^. The model statistics were validated using MolProbity^57^. Structural figures were prepared in Chimerax^58^ and PyMOL (https://pymol.org/2/). For the inactive LHCGR, the model of active LHCGR from the structure of Org43553/CG-LHCGR-Gs complex was to generate the initial template. The initial template was docked into the EM density map using UCSF Chimera and was subjected to flexible fitting using Rosetta^59^. The model was further rebuilt in coot and real-space refined in Rosetta and Phenix. The final refinement statistics were validated using the module “comprehensive validation (cryo-EM)” in Phenix. The final refinement statistics are provided in Extended data Table 1.

### Molecular dynamics simulation and data analysis

The cryo-EM structure of the CG-bound LHCGR-Gs complex was used to build the model of LHCGR with and without CG. Meanwhile, the cryo-EM structure of LHCGR-Gs structure with Org43553 and CG was applied to build the model of Org43553 bound system. Missing atoms and loops were added back in the most favored position without clashes using Modeller^60^. Using CHARMM-GUI^61, 62^, the models were inserted in POPC (palmitoyl-2-oleoyl-sn-glycero-3-phosphocholine) lipids to establish a CG-bound LHCGR-Gs complex system and a LHCGR-only system, and a CG-bound LHCGR-Gs complex system with Org43553, respectively. In each simulation system, the protein model and lipid bilayer were solvated in a periodic boundary condition box (13.8 nm × 13.8 nm× 22.4 nm) filled with TIP3P (transferable intermolecular potential 3 P) water molecules^63^ and 0.15 M KCl. On the basis of the CHARMM all-atom force field^64, 65^, 300- ns simulation was performed for each system using GROMACS^66, 67^. The system with Org43553 has five individual runs. The ligand parameters were derived from the CHARMM general force field (CGenFF) for small molecule drug design. No extra potential restraints were applied. All productions were performed in the NPT ensemble at a temperature of 303.15 K and a pressure of 1 atm. Temperature and pressure were controlled using the velocity-rescale thermostat^68^ and the Parrinello–Rahman barostat with isotropic coupling^69^, respectively. Equations of motion were integrated with a 2 fs time step, and the LINCS algorithm was used to constrain bonds lengths^70^. Nonbonded pairlists were generated every 10 steps using a distance cutoff of 1.4 nm. A cutoff of 1.2 nm was used for Lennard-Jones (excluding scaled 1–4) interactions, which were smoothly switched off between 1 and 1.2 nm. Electrostatic interactions were computed using the Particle-Mesh-Ewald algorithm^71^ with a real-space cutoff of 1.2 nm. Trajectory analysis was performed using GROMACS.

## Acknowledgments

The cryo-EM data were collected at the Cryo-Electron Microscopy Research Center, Shanghai Institute of Materia Medica (SIMM) and Shuimu BioSciences Ltd (Beijing China). The authors thank the staff at the SIMM Cryo-Electron Microscopy Research Center and the Shuimu BioSciences Ltd. for their technical support. This work was partially supported by Ministry of Science and Technology (China) grants (2018YFA0507002 to H.E.X.); Shanghai Municipal Science and Technology Major Project (2019SHZDZX02 to H.E.X.); CAS Strategic Priority Research Program (XDB08020303 to H.E.X.); the National Natural Science Foundation of China (31770796 to Y.J.); the National Science & Technology Major Project “Key New Drug Creation and Manufacturing Program” (2018ZX09711002 to Y.J.); Wellcome Trust grant 209407/Z/17/Z; National Key R&D Program of China (2020YFC0861000 to S.Z.); CAMS Innovation Fund for Medical Sciences (2020-I2M-CoV19-001 to S.Z.); Tsinghua University-Peking University Center for Life Sciences (045-160321001 to S. Z.); National Science & Technology Major Project “Key New Drug Creation and Manufacturing Program” of China (2018ZX09711002 to H.J.); Science and Technology Commission of Shanghai Municipal (20431900100 to H.J.); Jack Ma Foundation (2020-CMKYGG-05 to H.J.); Shanghai Sailing Program (19YF1457600 to Q.F.L.).

## Author contributions

J.D. designed the expression constructs, purified the LHCGR proteins, prepared the final samples for negative stain and data collection toward the structures, and conducted functional studies, and participated in figure and manuscript preparation; P.X. performed cryo-EM grid preparation, cryo-EM data collection, map calculations, model building, and participated in figure preparation; X.C. and X.H. performed MD stimulation analysis and participated in figure preparation; C.M. processed the datasets of the Org43553 bound LHCGR and the inactive LHCGR, and built the model for the inactive structure; T.C. helped build and refine the structure model; J.S and E.L.Y synthesized Org43553 and Compound 26; W.Y. designed G protein constructs; X.L and S.Z. coordinated EM data collection; Q.L. performed detergent and lipid screen experiments; H.J. supervised and coordinated MD experiments; Y. Z. supervised C.M. in data processing and figure preparation; Y.J. supervised the studies, performed the structural analysis, and participated in manuscript preparation; H.E.X. conceived and supervised the project, analyzed the structures, and wrote the manuscript with inputs from all authors.

## Data availability

The density maps and structure coordinates have been deposited to the Electron Microscopy Database (EMDB) and the Protein Data Bank (PDB) with accession number of EMD-31596, PDB ID 7FIG for the CG-LHCGR(S277I)-Gs complex; EMD-31597 and 7FIH for the CG-Org43553-LHCGR(S277I)-Gs complex; EMD-31598 and 7FII for the CG-LHCGR(WT)-Gs complex; EMD-31599 and 7FIJ for the LHCGR (inactive) structure.

---

**Extended Data Fig. 1.**
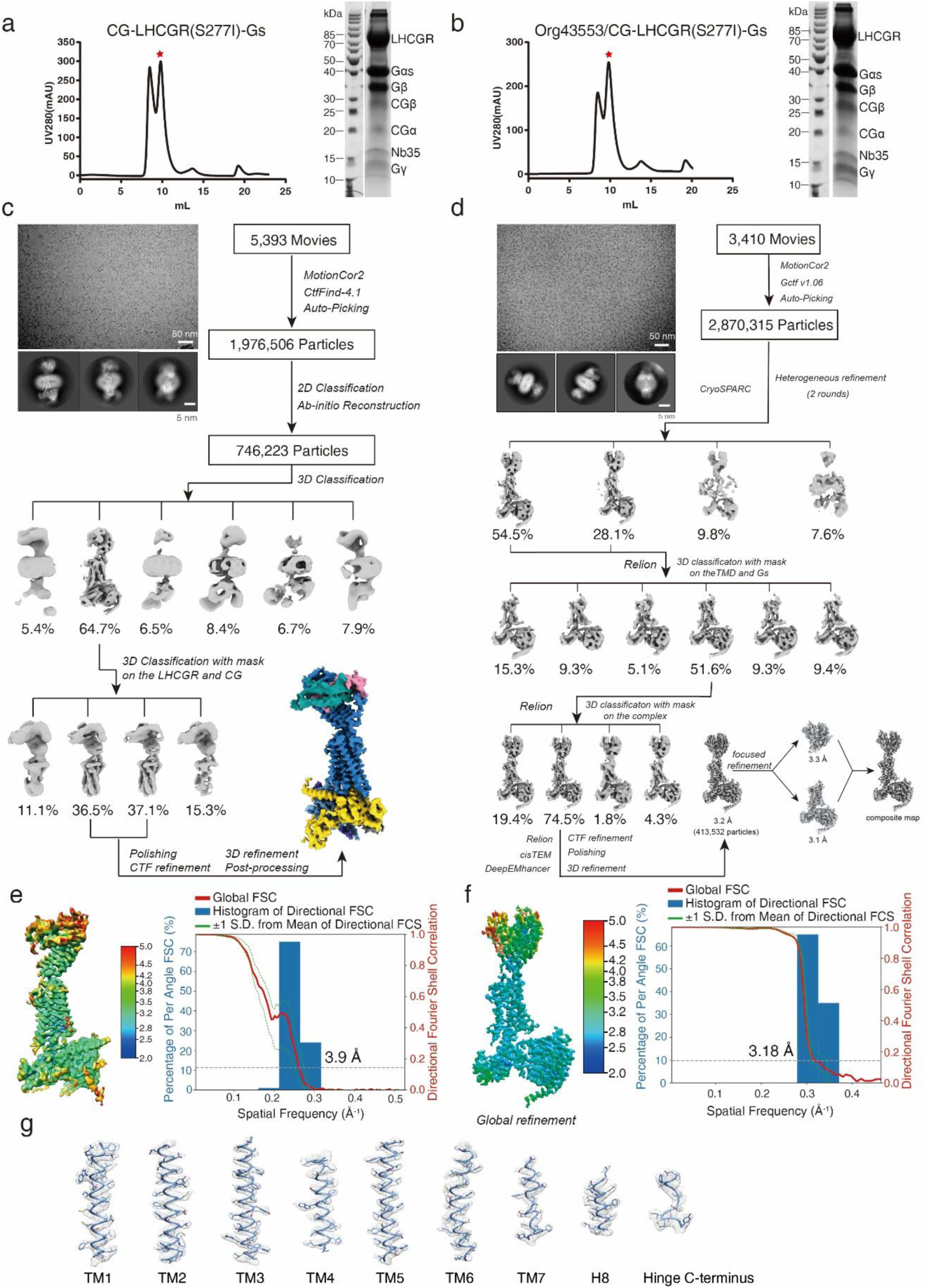
Cryo-EM images and single-particle reconstruction of the CG-LHCGR-Gs and Org43553/CG-LHCGR-Gs complexes, as well as EM maps for the Org43553/CG-LHCGR-Gs complex. **a**,**b** Size-exclusion chromatography elution profiles and SDS-PAGEs of the CG-LHCGR-Gs complex (left panel) and Org43553/CG-LHCGR-Gs complex (right panel). Red stars indicate the monomer peaks of the two complexes. **c,d** Cryo-EM micrograph, reference-free 2D class averages, and flowchart of cryo-EM data analysis of the CG-LHCGR-Gs complex (**c**) and Org43553/CG-LHCGR-Gs complex (**d**). **e,f** Cryo-EM maps of the CG-LHCGR-Gs (**e**) and Org43553/CG-LHCGR-Gs (**f**) complexes colored by local resolutions from 2.0 Å (blue) to 5.0 Å (red).The “Gold-standard” Fourier shell correlation (FSC) curves indicate that the overall resolution of the electron density map of the CG-LHCGR-Gs complex is 3.9 Å (**e**) and the Org43553/CG-LHCGR-Gs complex is 3.18 Å (**f**). **g**, Cryo-EM density map and model of the Org43553/CG-LHCGR-Gs complex. The regions of the cryo-EM density map with all transmembrane helices, H8 and hinge C-terminus are shown.

**Extended Data Fig. 2.**
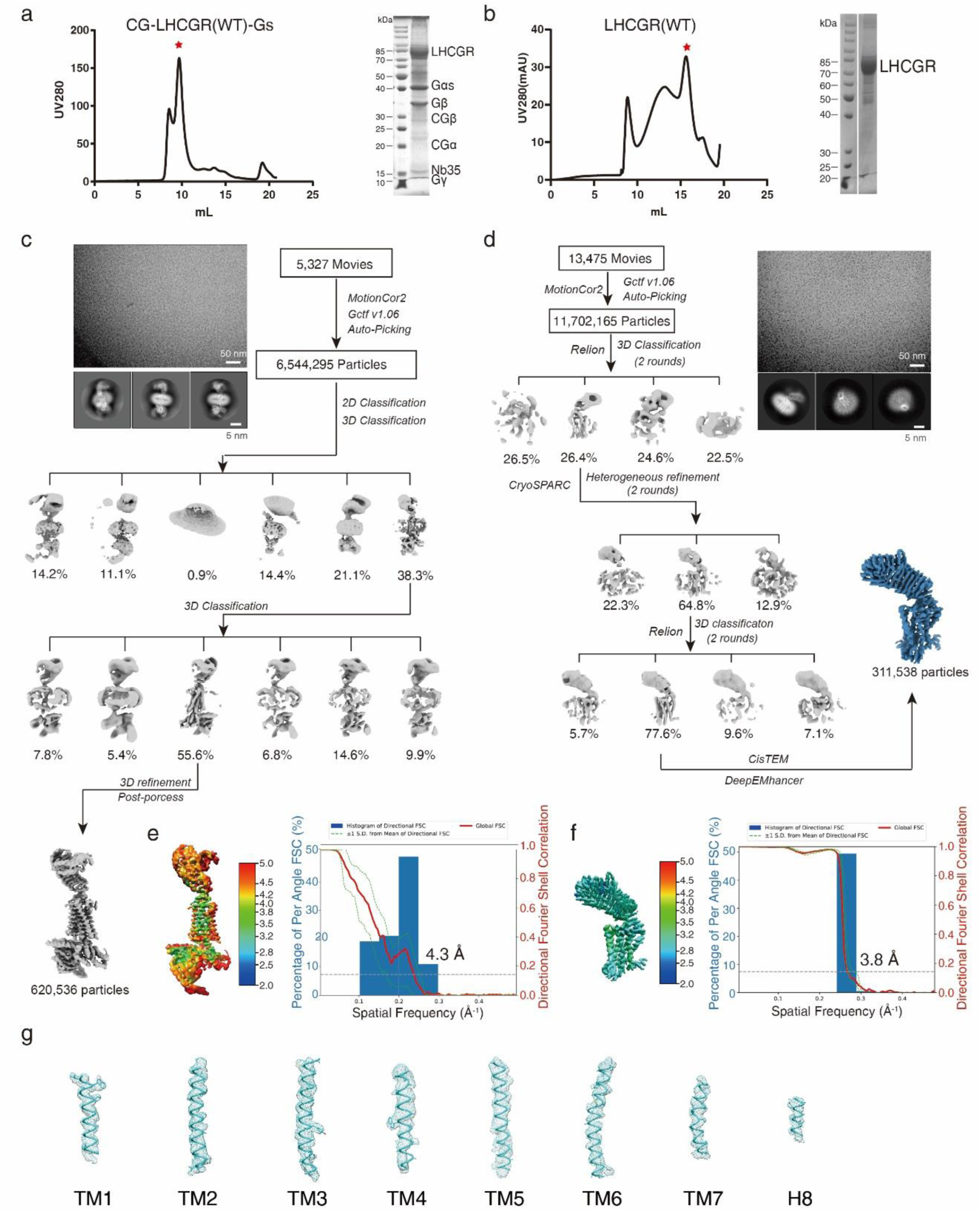
Cryo-EM images and single-particle reconstruction of the Org43553/CG-LHCGR(WT)-Gs complex and inactive LHCGR, as well as EM maps for the inactive LHCGR structure. **a**,**b** Size-exclusion chromatography elution profiles and SDS-PAGEs of the Org43553/CG-LHCGR(WT)-Gs complex (left panel) and inactive LHCGR (right panel). Red stars indicate the monomer peaks of the two proteins. **c,d** Cryo-EM micrograph, reference-free 2D class averages, and flowchart of cryo-EM data analysis of the Org43553/CG-LHCGR(WT)-Gs complex (**c**) and inactive LHCGR (**d**). **e,f** Cryo-EM maps of the Org43553/CG-LHCGR(WT)-Gs complex (**e**) and inactive LHCGR (**f**) colored by local resolutions from 2.0 Å (blue) to 5.0 Å (red).The “Gold-standard” Fourier shell correlation (FSC) curves indicate that the overall resolution of the electron density map of the Org43553/CG-LHCGR(WT)-Gs complex is 4.3 Å (**e**) and inactive LHCGR is 3.8 Å (**f**) . **g**, Cryo-EM density map and model of the inactive LHCGR. The regions of the cryo-EM density map with all transmembrane helices and H8 are shown.

**Extended Data Fig. 3.**
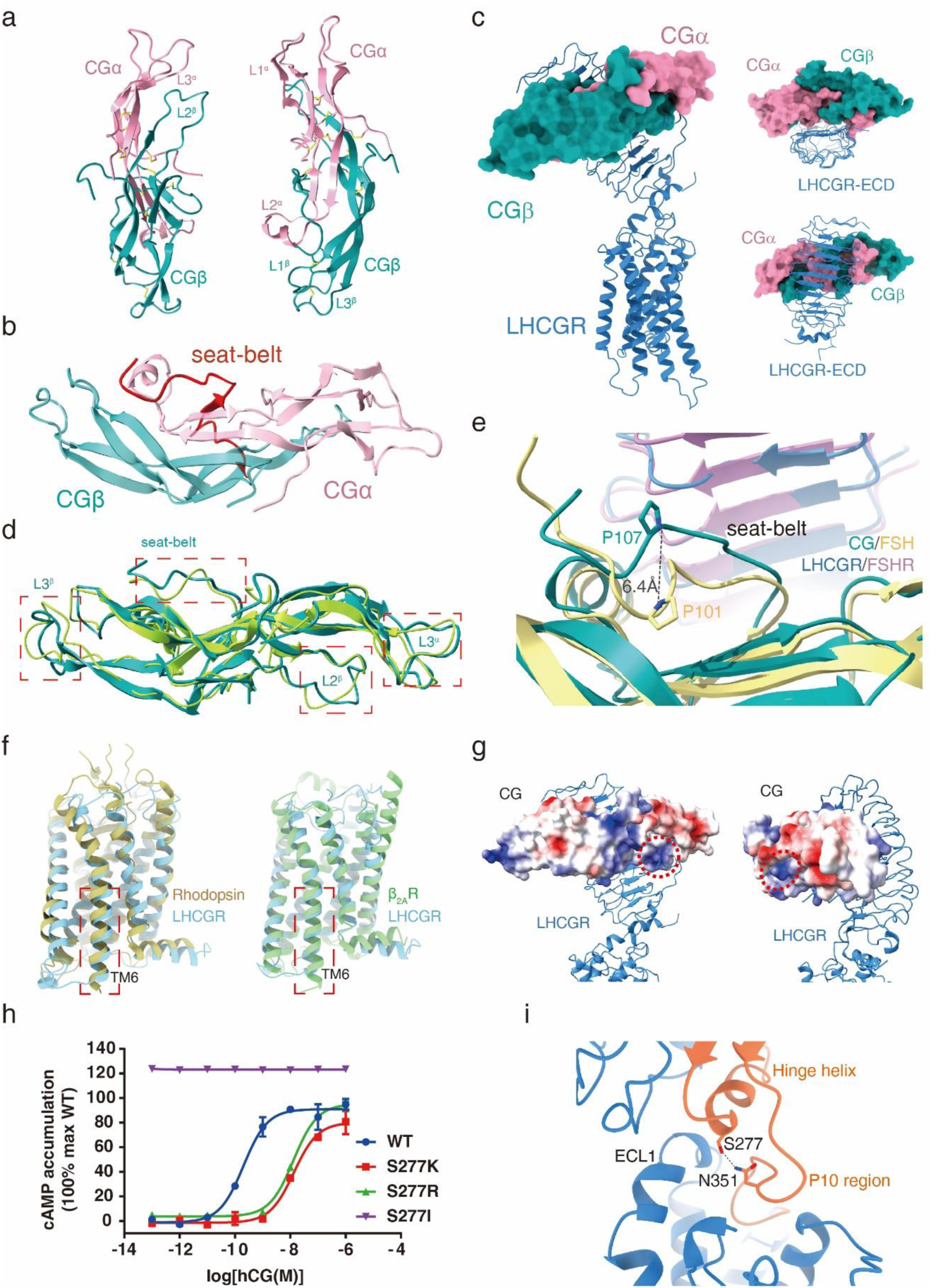
Structural features of LHCGR. **a**, Ribbon presentation of CGα and CGβ subunits stabilized by cysteine-knots. Disulfide bonds are shown as yellow sticks. **b**, Ribbon presentation of CGα and CGβ subunits. The C-terminus of the CGβ subunit, known as “seat-belt” highlighted in red, circulates the CGα subunit. **c**, A “hand-clasp” binding fashion of CG to LHCGR from different views. CGα and CGβ subunits are shown in a surface presentation, while LHCGR is displayed as a ribbon**. d**, Structural comparison of free human CG (PDB code: 1HCN, green-yellow) with receptor-bound CG (light sea green), the conformational changes in four segments are highlighted in red rectangle. **e**, Structural comparison of “seat-belt” between CG-LHCGR and FSH-FSHR(PDB code: 1XWD). The residue P107 in CGβ and P101 in FSHβ are shown as sticks. **f**, Conformational comparison of TM6 of inactive LHCGR with inactive rhodopsin (PDB code:1L9H, left panel) and inactive β_2_AR (PDB code: 5JQH, right panel). **g**, Electrostatic potential surface of CG. The positively charged pockets that interact with the hinge loop are highlighted in red circles. **h**, Concentration-response curves for point mutants at S277. Experiments were performed in triplicate, and the representative concentration-response curves were presented. **i**, The putative interaction between S277 in the hinge helix and N351 in the P10 region based on the model of Org43553/CG-LHCGR-Gs complex.

**Extended Data Fig. 4.**
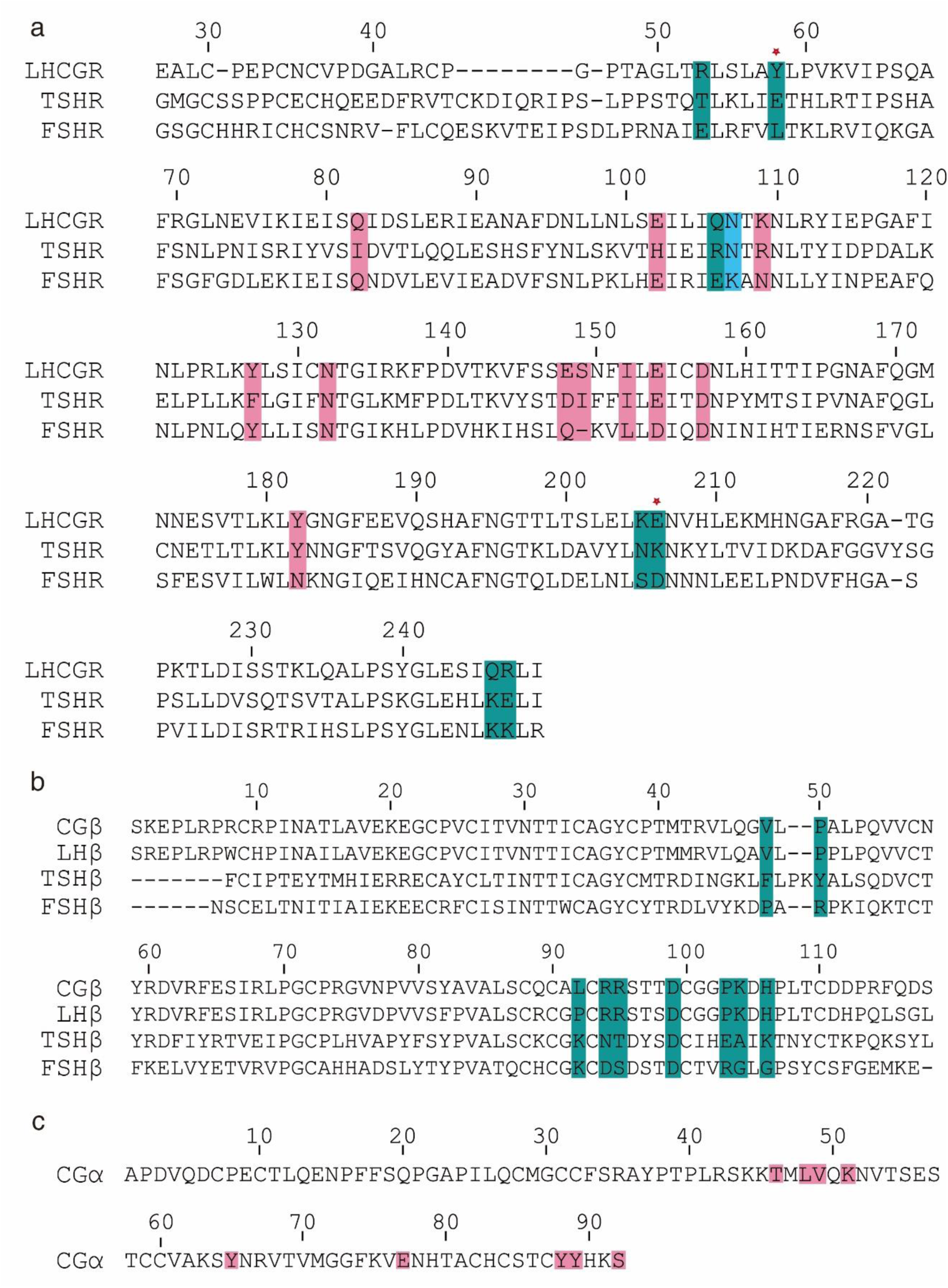
Sequence alignment of glycoprotein hormones and related receptors. **a**, Sequences alignment of human FSHR, LHR and TSHR in the region of the hormone-binding domain. Residues only interact with CGβ (light sea green), CGα (pink), and both CGα and CGβ subunits (light blue) are highlighted. Residues that determine LHCGR specificity were labeled with asterisks. **b**, Sequences alignment of human CG, LH, TSH and FSH β-subunit. CGβ residues interacted with LHCGR are highlighted in light sea green. **c**, The α-subunit sequence of glycoprotein hormones. CGα residues interacted with LHCGR are highlighted in pink.

**Extended Data Fig. 5.**
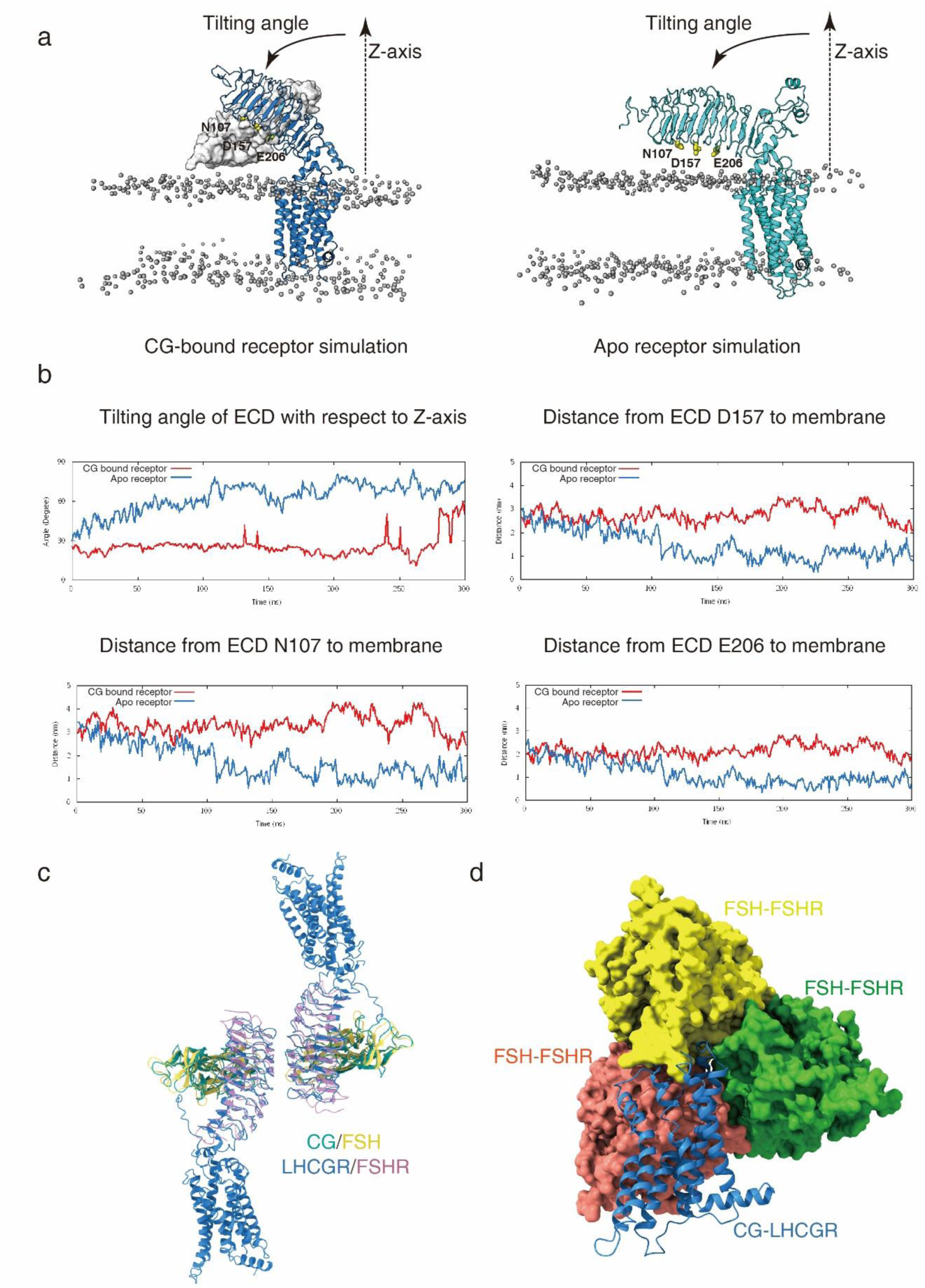
Molecular dynamics simulations showing hormone-induced receptor activation. **a**, Representative snaphots from the CG-bound receptor simulation and inactive receptor simulation, respectively. The receptor is shown as cartoon, the CG is shown as surface, the phosphate groups of the lipid membrane are shown as spheres, three ECD residues N107, D157 and E206 are shown as sticks. **b**, Changes of ECD orientations observed in the different simulations as a function of time. Tilting angle of ECD with respect to the Z-axis (membrane normal) and the minimal distance from an ECD residue (D157, N107, or E206) to the membrane. **c**, Structural alignment of the CG-LHCGR complex onto the FSH-FSHR-ECD dimer (PDB code: 1WXD, left panel) shows that the two TMD from each complex is pointed in the opposite direction and cannot be on the same membrane layer. **d**, Structural alignment of the CG-LHCGR complex onto the FSH-FSHR-ECD trimer (PDB code: 4AY9, right panel) shows that the TMD from the CG-LHCGR complex will clash with the neighboring ECD in the trimeric arrangement. The trimeric FSH-FSHR-ECD was shown in surface presentation, with each monomeric complex in a single color (red, yellow, green). The CG-LHCGR is shown as a ribbon and is aligned onto the yellow ECD complex but is clashed with the red ECD complex.

**Extended Figure 6.**
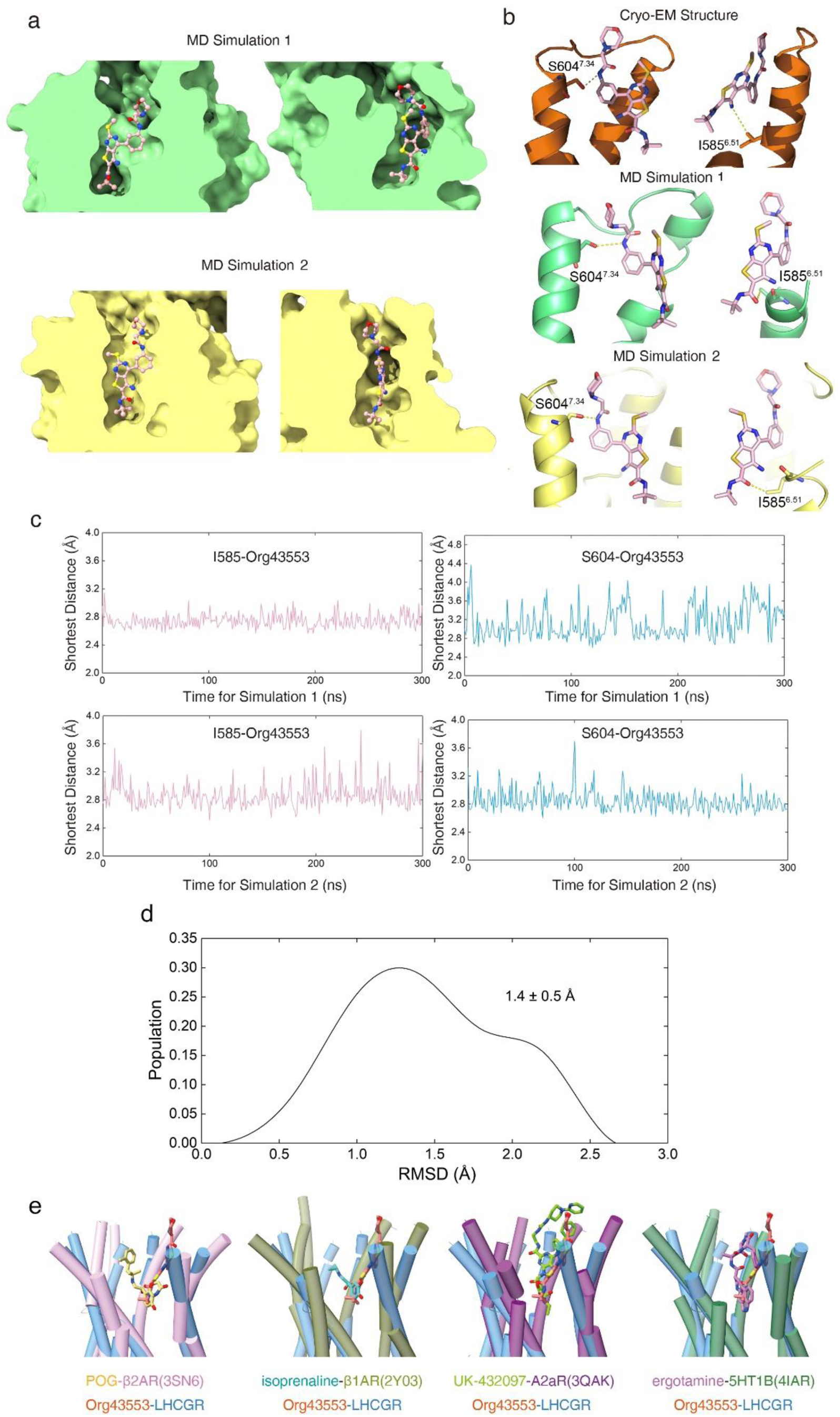
The binding pocket of Org43553 in the top half of the TMD. **a,** Two views of the Org43553 binding pocket in the representative structures of MD Simulation 1 and 2. Org43553 is shown as sticks, while LHCGR is depicted by surface. **b**, The interactions between I585, S604 and Org43553 in the cryo-EM structure and the representative structures of simulation 1 and simulation 2. Org43553 is shown as sticks, while LHCGR is shown as cartoon. **c**, The time-course curve of the shortest distance between the heavy atoms of I585, S604 and Org43553 during simulation 1 and 2. **d**, The distribution of the RMSD from the cryo-EM ligand pose in simulations of Org43553- bound LHCGR system. **e**, Comparison of the Org43553-binding pocket with other agonist-binding pockets of class A GPCRs. β_2_AR (PDB code: 3SN6); β_1_AR (PDB code: 2Y03); A_2a_R (PDB code: 3QAK); 5HT_1B_ (PDB code: 4IAR); Org43553, tomato; β_2_AR-agonist, yellow; β_1_AR-agonist, cyan; A_2a_R-agonist, lime green; 5HT_1B_-agonist, magenta.

**Extended Data Fig. 7.**
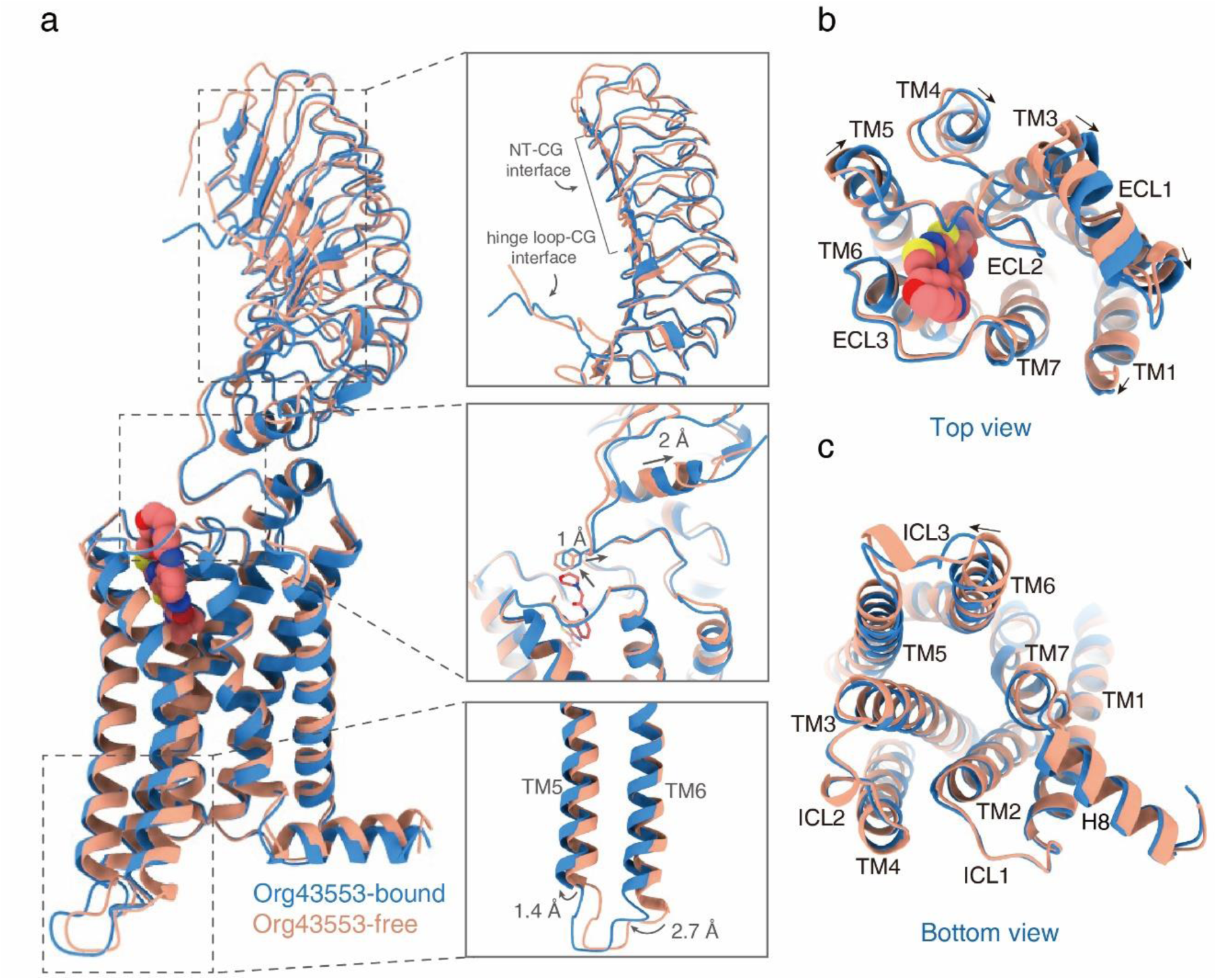
Comparison of the Org43553-bound LHCGR structure with the Org43553-free LHCGR structure. **a,** Superposition of the Org43553-bound LHCGR and Org43553-free LHCGR. **b**, Snapshots of the comparison from the top view. **c**, Snapshots of the comparison from the bottom view. Org43553-bound receptor is colored with blue, Org43553-free receptor is colored with light pink.

**Extended Data Fig. 8.**
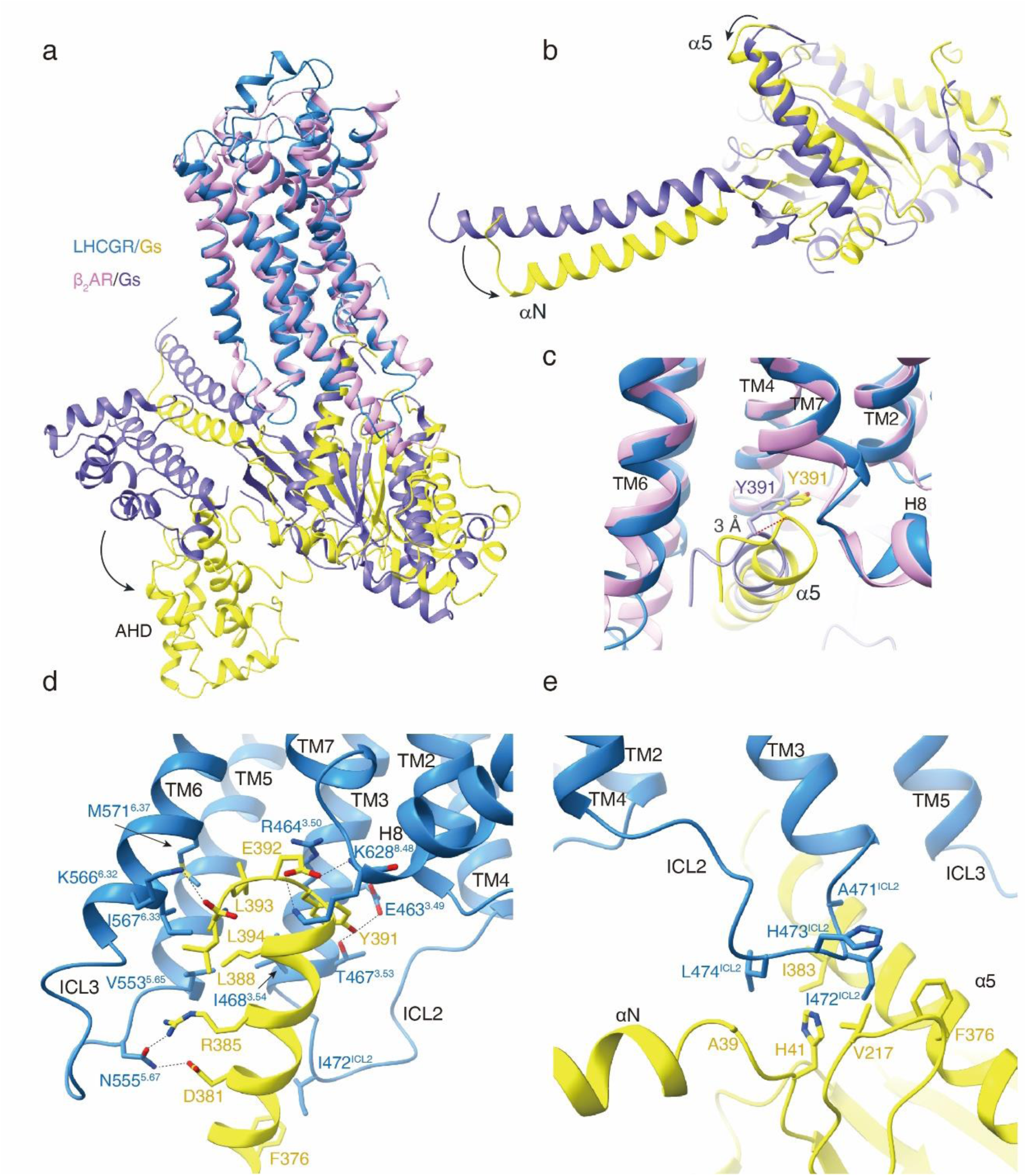
Gs-protein coupling of LHCGR. **a**, A conformational comparison of receptor helical bundle and Gαs between CG-LHCGR-Gs and β_2_AR-Gs complexes. **b**, A comparison of the rotation of Gαs between CG-LHCGR-Gs and β_2_AR- Gs complexes. **c**, The α5 helix of Gαs in LHCGR-Gs inserts 3 Å deeper into the cytoplasmic TMD bundle relative to that in β_2_AR-Gs complex. **d**,**e** Detailed interactions of receptor TM helices (TM3, TM5, TM6, TM7, and Helix 8) with α5 helix of Gαs (**d**), as well as ICL2 of the receptor with αN and α5 helix of Gαs (**e**). LHCGR and Gs are colored in blue and yellow, while β_2_AR and Gs in hot pink and slate, respectively.

**Extended Data Fig. 9.**
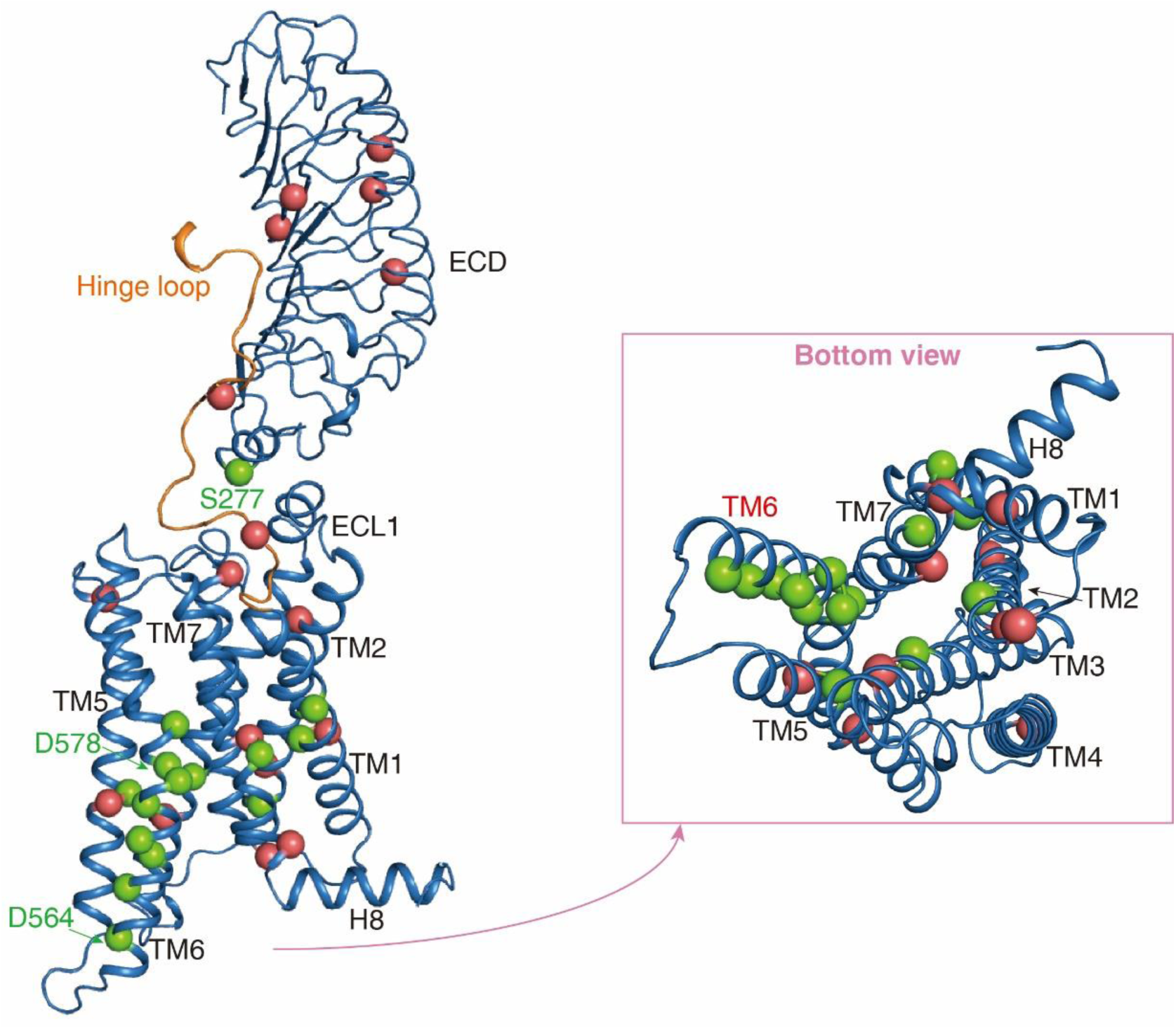
Missense mutations in LHCGR. The inactive mutations are highlighted in red spheres, and the constitutively active mutations are highlighted in green spheres.

**Extended Data Fig. 10.**
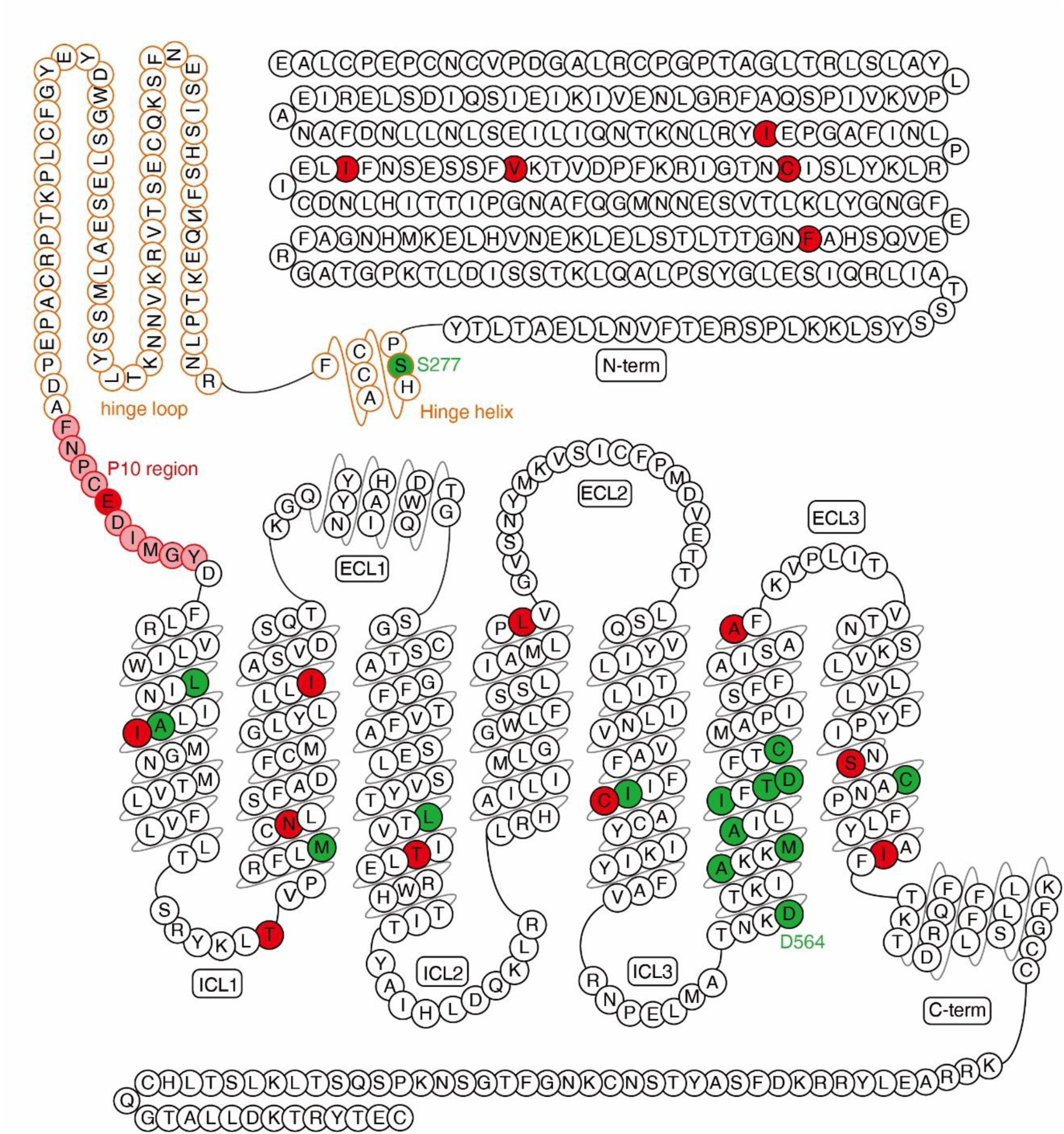
Snake model of the LHCGR. The P10 region is indicated in pink cycles. The inactive mutations are highlighted in red cycles, and the constitutively active mutations are highlighted in green cycles.

**Extended data Table 1:**
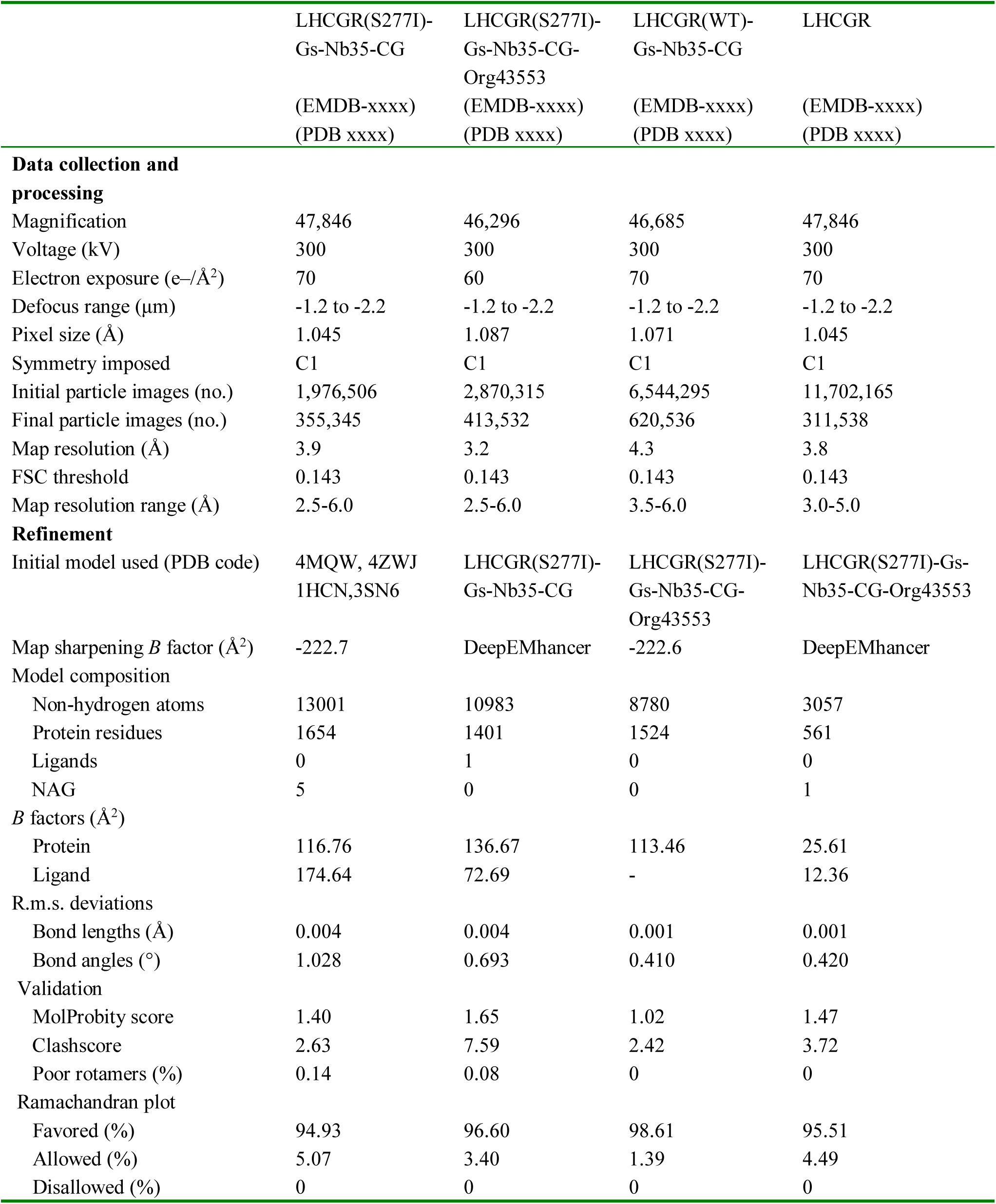
Cryo-EM data collection, refinement and validation statistics.

**Extended data Table 2:**
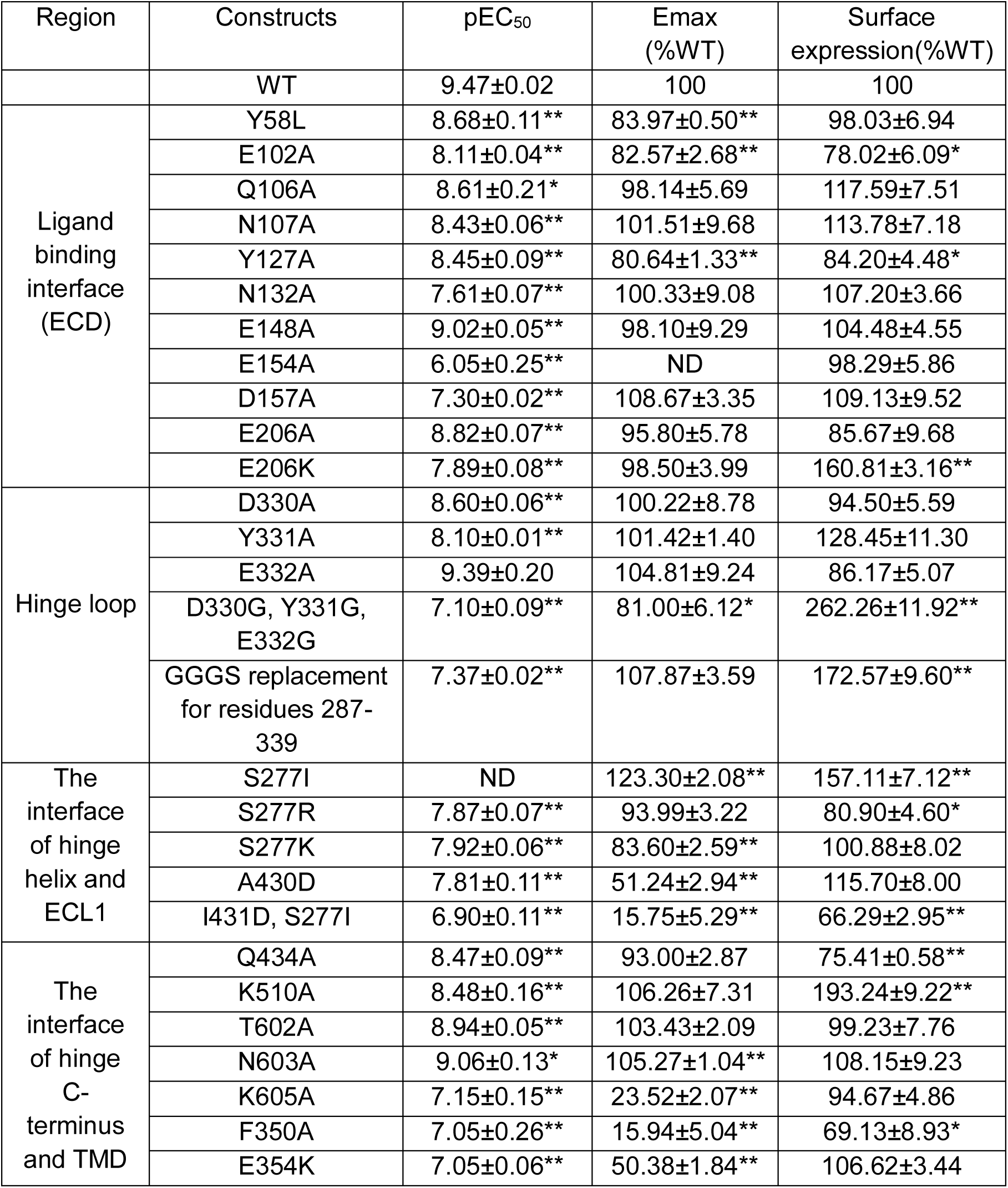
CG-induced activation on wild type and LHCGR with site-directed mutations. Data represent mean pEC50 (pEC50 ± SEM), Emax (Emax ± SEM). Experiments were performed in triplicate. The Emax and surface expression of LHCGR mutants were normalized to WT, which was set to 100. *P<0.05, **P<0.01 versus WT.

**Extended data Table 3:** Sequence alignment of residues from the four conserved motifs in class A GPCRs. Receptors are all from class A GPCR family.

|  | DRY motif |  |  | PIF motif |  |  | Toggle switch | NPXXY motif |  |  |
| --- | --- | --- | --- | --- | --- | --- | --- | --- | --- | --- |
|  | 3.49 | 3.50 | 3.51 | 5.50 | 3.40 | 6.44 | 6.48 | 7.49 | 7.50 | 7.53 |
| [Human]LHCGR | E | R | W | A | V | D | M | N | P | Y |
| [Human]FSHR | E | R | W | A | V | D | M | N | P | Y |
| [Human]TSHR | E | R | W | A | V | D | M | N | P | Y |
| [Human] beta2-adrenoceptor | D | R | Y | P | I | F | W | N | P | Y |
| [Human] A2A receptor | D | R | Y | P | I | F | W | N | P | Y |
| [Human] A1 receptor | D | R | Y | P | I | F | W | N | P | Y |
| [Human] μ receptor | D | R | Y | P | I | F | W | N | P | Y |
| [Human] H1 receptor | D | R | Y | P | I | F | W | N | P | Y |
| [Human] 5-HT1A receptor | D | R | Y | P | I | F | W | N | P | Y |
| [Human] D2 receptor | D | R | Y | P | I | F | W | N | P | Y |
| [Human] CB2 receptor | D | R | Y | L | V | L | W | N | P | Y |
| [Human] CXCR4 | D | R | Y | P | V | F | W | N | P | Y |
| [Human] CCR1 | D | R | Y | P | I | F | W | N | P | Y |
| [Human] M1 receptor | D | R | Y | P | V | F | W | N | P | Y |
| [Human] Y1 receptor | E | R | H | P | I | F | W | N | P | Y |
| [Human] CCK1 receptor | E | R | Y | P | T | F | W | N | P | Y |
| [Human] ghrelin receptor | E | R | Y | P | V | F | W | N | P | Y |
| [Human] MC1 receptor | D | R | Y | M | L | F | W | D | P | Y |
| [Human] MCH1 receptor | D | R | Y | P | T | F | W | N | P | Y |
| [Human] NTS1 receptor | E | R | Y | P | A | F | W | N | P | Y |
| [Human] PAR1 | D | R | F | P | I | F | F | D | P | Y |
| [Human] ETB receptor | D | R | Y | P | V | F | W | N | P | Y |
| [Human] V2 receptor | D | R | H | P | S | Y | W | N | P | Y |
| [Human] OX1 receptor | D | R | W | P | V | F | Y | N | P | Y |
| [Human] GnRH1 receptor | D | R | S | P | A | F | W | D | P | Y |
| [Human] kisspeptin receptor | D | R | W | P | C | F | W | N | P | Y |
| [Human] AT1 receptor | D | R | Y | P | V | F | W | N | P | Y |
| [Human] GAL1 receptor | D | R | Y | P | I | F | W | N | P | Y |
| [Human] NPBW1 receptor | D | R | Y | P | L | C | W | N | P | Y |
| [Human] motilin receptor | E | R | Y | P | L | F | W | N | P | Y |
| [Human] NK2 receptor | D | R | Y | P | I | F | W | N | P | Y |
| [Human] SST1 receptor | D | R | Y | P | I | F | W | N | P | Y |
| [Human] FPR2/ALX | D | R | C | P | V | F | W | N | P | Y |
| [Human] NK2 receptor | D | R | Y | P | I | F | W | N | P | Y |
| [Human] TA1 receptor | D | R | Y | P | I | F | W | N | P | Y |
| [Human] EP1 receptor | E | R | C | L | L | S | W | D | P | Y |
| [Human] apelin receptor | D | R | Y | P | V | F | W | N | P | Y |
| [Human] NPFF1 receptor | E | R | F | P | V | F | W | N | P | Y |

